# Infants’ cortex undergoes microstructural growth coupled with myelination

**DOI:** 10.1101/2021.03.16.435703

**Authors:** Vaidehi S. Natu, Mona Rosenke, Hua Wu, Francesca R. Querdasi, Holly Kular, Nancy Lopez-Alvarez, Mareike Grotheer, Shai Berman, Aviv A. Mezer, Kalanit Grill-Spector

## Abstract

Development of cortical tissue during infancy is critical for the emergence of typical brain functions in cortex. However, how cortical microstructure develops during infancy remains unknown. We measured the longitudinal development of cortex from newborns to six-months-old infants using multimodal quantitative imaging of cortical microstructure. Here we show that infants’ cortex undergoes profound microstructural tissue growth during the first six months of human life. Comparison of postnatal to prenatal transcriptomic gene expression data demonstrates that myelination and synaptic processes are dominant contributors to this postnatal microstructural tissue growth. Using visual cortex as a model system, we find hierarchical microstructural growth: higher-level visual areas have less mature tissue at birth than earlier visual areas but grow at faster rates. This overturns the prevailing view that visual areas that are most mature at birth develop fastest. Together, *in vivo*, longitudinal, and quantitative measurements, which we validated with *ex vivo* transcriptomic data, shed new light on the rate, sequence, and specific biological mechanisms of developing cortical systems. Importantly, our findings propose a new hypothesis that cortical myelination is a key factor in cortical development during early infancy, which has significant implications for diagnosis of neurodevelopmental disorders and delays in infants.

## Introduction

Development of the neuroarchitecture during early infancy is critical for the maturation of key sensory and cognitive functions and has lifelong consequences. Primary sensory-motor cortices are more developed in infants than the prefrontal cortex^1–7^, which is involved in complex cognitive functions. However, the rate, sequence, and microstructural mechanisms underlying cortical tissue development in human infants during the first six months of life are unknown.

Present understanding of microstructural development is gleaned from histological investigations of a handful of sensory-motor and prefrontal regions in humans^2,3^ and non-human primates^8–10^. These studies suggest that during development, cortex grows synapses^2,9,11^, dendrites^10,12^, axons^10,12^, and myelin^3^, but also prunes irrelevant connections and synapses^2,10,12,13^. However, there is an intense debate regarding the relative effects of microstructural growth and pruning and if they vary across cortical regions.

To fill these glaring gaps in knowledge, we leveraged recent advancements in quantitative magnetic resonance imaging (qMRI)^14,15^ and diffusion MRI (dMRI)^16^ to develop novel *in vivo* methodologies that are optimized for the infant brain. Research in humans is critical, as compared to other species, humans have protracted brain development and unique brain structure^3^. Quantitative measurements of proton relaxation time (T_1_, which depends on the physiochemical tissue environment) from qMRI and mean diffusivity (MD, which depends on the density and structure of tissue through which water diffuses) from dMRI enable quantifying and longitudinally measuring the amount of brain tissue within a voxel (3D pixel in an MRI image, 1-2mm on a side) related to the neuropil^17^ and myelin^18^.

Critically, different from standard MRI, these measurements have metric values with units, and can disambiguate developmental hypotheses as T_1_ and MD are lower in tissue with denser microstructure^15,17,18^. Thus, we predicted that if cortical microstructure proliferates, T_1_ and MD will decrease during infancy, but if microstructure is pruned, T_1_ and MD will increase. We tested these hypotheses in (i) primary sensory-motor areas to relate our measurements to prior *ex vivo* studies, and (ii) across two visual processing hierarchies^19^, which offer an exciting opportunity to investigate the sequence and rate of microstructural development across an entire cortical system for the first time.

Anatomical MRI, qMRI, and dMRI data were collected in 13 full-term infants (6 female) who were scanned during normal sleep at 0 months [8-37 days], 3 months [78-106 days], and 6 months [167-195 days] (10 infants per timepoint, 7 infants scanned longitudinally at all timepoints, **Supplementary Fig. S1** and Methods). For quality assurance, we (i) monitored in real-time each infant’s motion via an infrared camera, (ii) assessed the quality of brain images immediately after acquisition, and (iii) repeated scans with motion artifacts. From anatomical MRIs, we generated the cortical surface for each infant and timepoint (**Supplementary Fig. S1**). Cortical surface reconstruction enabled achieving the most precise measurements by (i) analyzing T_1_ and MD data in each infant’s native cortical space, and (ii) using cortical-based alignment to delineate known cortical areas^20,21^ in each infant’s brain.

### Primary sensory cortices undergo exuberant microstructural tissue growth during the first 6 months of life

Longitudinal cortical T_1_ maps reveal that T_1_ decreases from newborns to 6 months-olds. This decrease is heterogenous across cortex (example infant: **Fig. 1A;** all infants: **Supplementary Fig. S2**). E.g., at 3 months, occipital cortex and the central sulcus have lower T_1_ (black arrows in **Fig. 1A**) than parietal and frontal cortices (red arrows in **Figs. 1A, Supplementary Fig. S2**) even as the entire cortex in 3 months-olds has lower T_1_ than in newborns.

**Fig 1.**
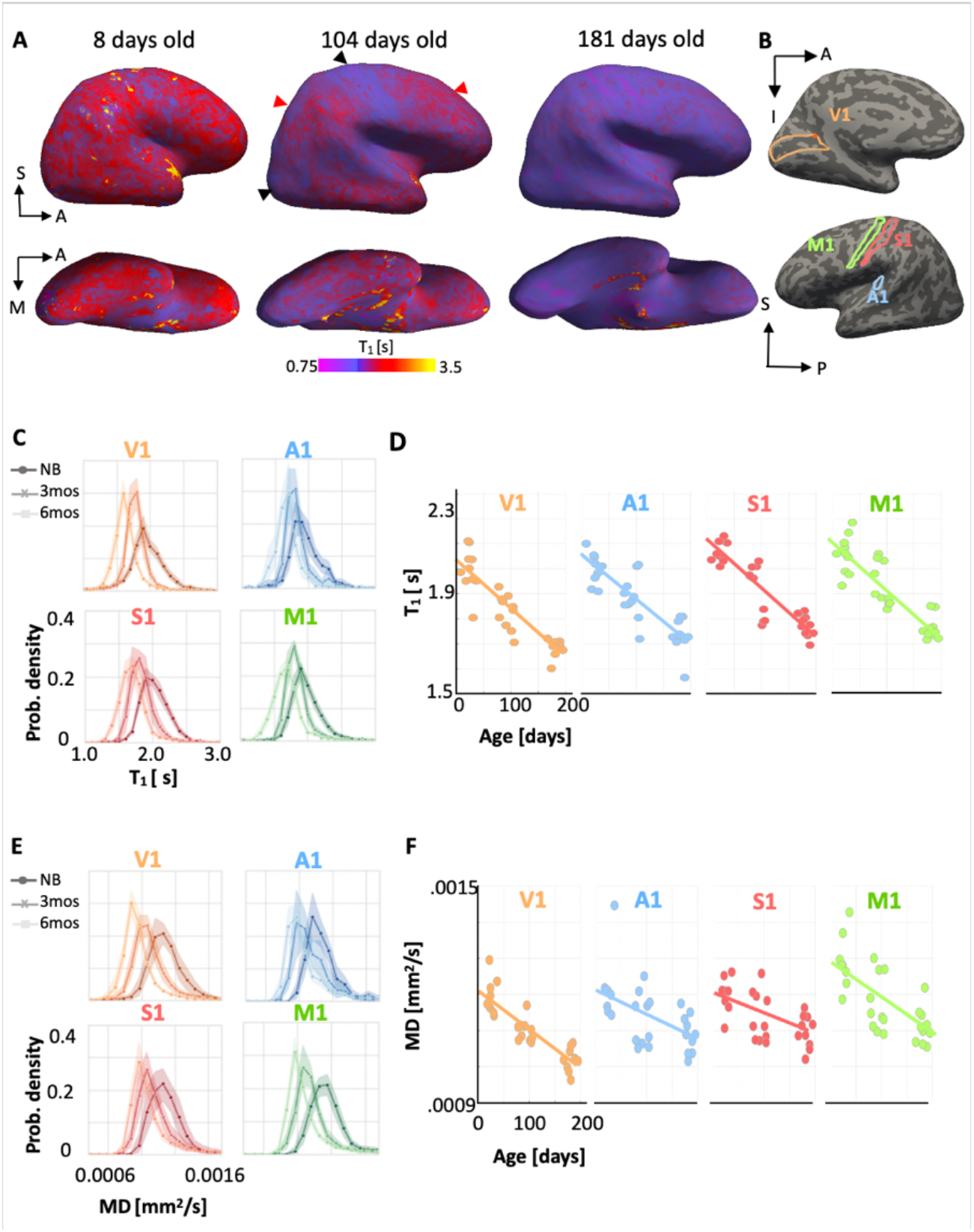
Primary sensory cortices are not fully developed at birth but show extensive microstructural tissue growth during the first 6 months of life. (A) Right hemisphere sagittal (top) and ventral-temporal (bottom) T_1_ maps [in seconds] displayed on an inflated cortical surface of an example baby across time. *Left to right:* cortical T_1_ at 8 days (newborn), 104 days (~ 3 months), and 181 days (~ 6 months) of age (*red/yellow:* higher T_1_; *purple*: lower T_1_). (B) Primary sensory-motor areas ^21^: V1, A1, M1 and S1 shown on the cortical surface of this baby. (C) T_1_ distributions across voxels of each area show a leftward shift from newborns (darker colors) to 6 month-olds (lighter colors). *Solid lines*: mean distribution; *Shaded region*: standard error of the mean (SE) across 10 infants at each timepoint. *NB:* newborns; *3 mo:* 3 month-olds; *6 mo:* 6 month-olds. *Darker colors* indicate younger infants. (D) T_1_ linearly decreases with age in primary sensory-motor areas. *Each dot*: mean T_1_ per area per infant. *Line:* Linear mixed model line fit. (E-F) Same as in C-D for mean diffusivity (MD).

Next, we quantitatively measured T_1_ and MD in four primary sensory-motor areas^21^: visual (V1), auditory (A1), somatosensory (S1), and motor (M1) (**Fig. 1B**), which overlap the cortical expanse showing rapid development. We found a systematic decrease in the distribution of T_1_ values from birth to 6 months in all primary areas (**Fig. 1C**). Results also revealed a significant linear decrease in mean T_1_ across all areas (linear mixed model (LMM) slopes: −1.3 to −2 [ms/day], *P*s<10^−7^, **Supplementary Table S2**, all stats, right hemisphere: **Fig. 1D**, left hemisphere: **Supplementary Fig. S3**). Across all primary sensory-motor regions, mean T_1_ substantially decreased from 2.03s ± 0.07s (mean ± standard deviation (SD)) in newborns, to 1.87s ± 0.08s at 3 months, to 1.74s ± 0.06s at 6 months. Analysis of MD in these areas revealed similar significant, linear decreases from 0 to 6 months (LMM slopes: −9.36×10^−7^ to −1.01×10^−6^ [mm^2^/s/day], *P*s<0.001, **Supplementary Table S3**, all stats right hemisphere: **Figs. 1E, 1F**, left hemisphere: **Supplementary Fig. S4**).

### Hierarchical and heterogeneous development of cortical microstructure in visual streams

We next used visual cortex as a model system to investigate microstructural development as it is the best understood cortical system and contains well-defined hierarchical processing streams. The ventral visual stream^19^ is involved in visual recognition, and the dorsal stream^19^ involved in visually guided actions and localization. In each infant and timepoint, we identified nine visual areas in the dorsal stream (V1d to IPS3) and eight in the ventral stream (V1v to PHC2) by using cortex-based alignment to project the Wang atlas^20^ of retinotopic visual areas to each individual infant’s brain at each timepoint. Then, we measured T_1_ and MD in each infant’s visual areas and timepoints.

Results reveal that within the first 6 months of life, T_1_ decreases in visual cortex on average by 0.36s – 0.54s in both the ventral and dorsal streams regions (**Figs. 2A-B,D-E**). Linear mixed modeling (LMM) per area quantified this development, revealing two main findings. First, T_1_ significantly decreases in all dorsal (T_1_ change/slopes: −3 to −2 [ms/day], *Ps*<10^−7^, **Fig. 2B**) and ventral visual areas (slopes: −1.9 to −1.5 [ms/day], *P*s<10^−6^, **Fig. 2E, Supplementary Table S4**, all stats). Second, LMM estimates of T_1_ at birth showed a systematic increase in T_1_ at birth ascending the hierarchy of each processing stream. In the dorsal stream, T_1_ at birth increases from V1d [2.0s±0.028s] to IPS3 [2.29s±0.027s] (**Fig. 2C**-right hemisphere, **Supplementary Fig. S5**-left hemisphere). In the ventral stream, it increases from V1v [2.01s±0.026s] to PHC2 [2.21+.027s] (**Fig. 2F**-right hemisphere, **Supplementary Fig. S5**-left hemisphere).

**Fig 2.**
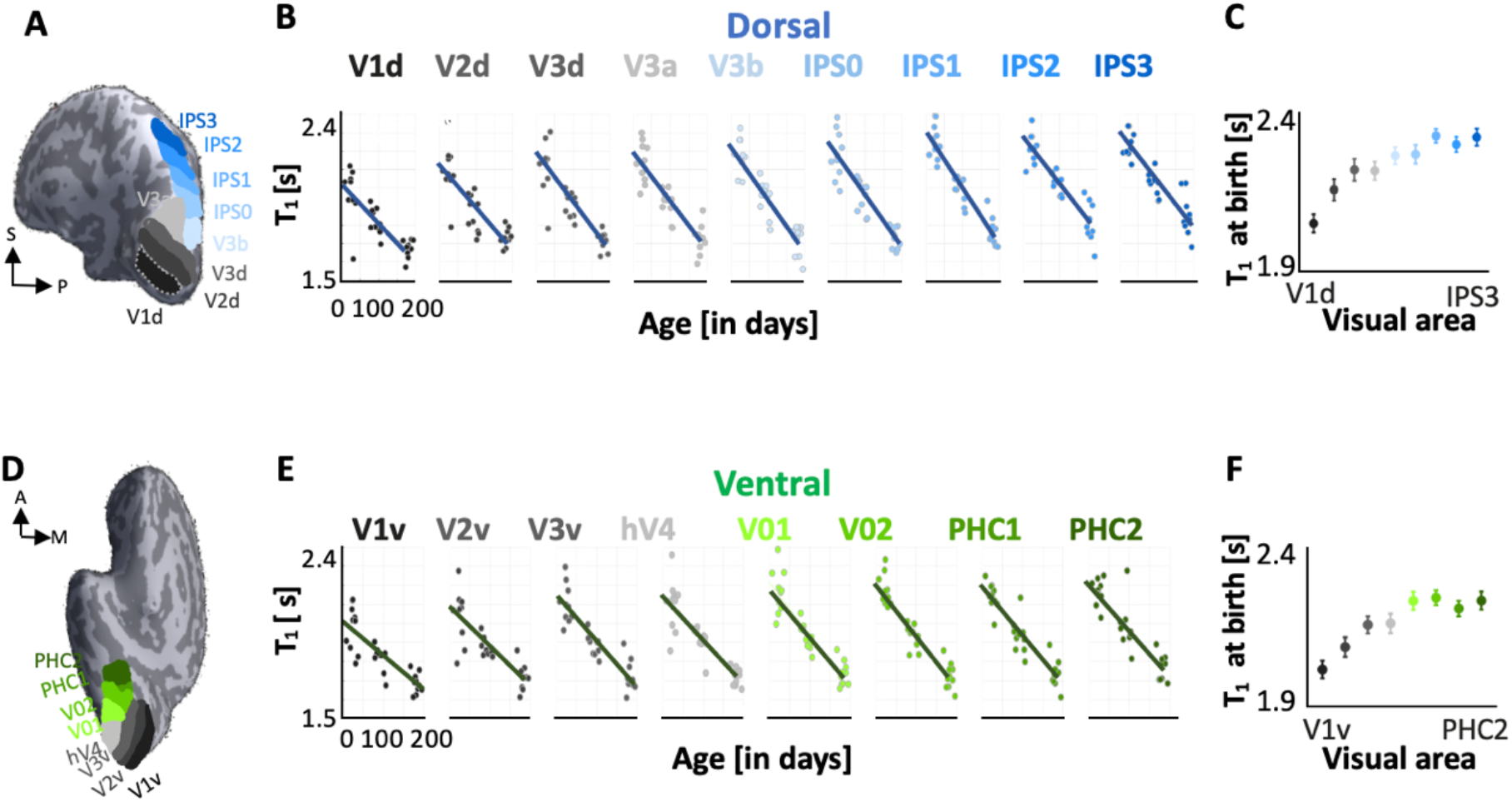
Hierarchical development of cortical microstructure in visual streams. (A,D) Inflated cortical surface of an example 6-month-old infant showing nine dorsal visual (A) and eight ventral visual (D) visual areas^20^. (B,E) T_1_ as a function of infant age in each visual area of dorsal (B) and ventral (E) visual processing streams. Color indicates the visual area, see A,D. *Each dot:* mean T_1_ per area per infant. Solid lines: Linear mixed model (LMM) estimates of T1 development per visual area. (C,F) LMM estimates of mean T_1_ at birth (LMM intercept) in each (D) dorsal and (F) ventral visual area. *Error bars:* standard error on estimates of intercepts. All data here are of the right hemisphere; left hemisphere data in **Supplementary Fig. S5**

In order to quantify the development in T_1_ across ROIs of each visual processing stream, we fitted LMMs relating the mean T_1_ of that area to participants’ age [in days] across all ROIs of the stream per hemisphere. Model comparison showed that an LMM which allows both intercepts and slopes to vary across ROIs best fit the data (Methods). This model revealed a significant main effect of ROIs across both streams and hemispheres (*ts* > 5.54, *Ps* < 8.045×10^−6^). These results suggest that there is hierarchical and heterogeneous development across both the ventral and dorsal visual processing hierarchies, as later visual areas have less mature tissue at birth than earlier visual areas but grow at faster rates (**Supplementary Table S4**-slopes). Similar results were observed with MD (*ts* > 2.25, *Ps* < 0.025) as MD significantly decreases from newborns to 6-month-olds and MD at birth systematically increases from V1 to later visual areas of each stream (**Supplementary Figs. S6, S7, Supplementary Table S5**).

Combined, these data reveal intriguing features of cortical development in the infant visual system. Ascending visual hierarchies from V1 to higher-level visual areas, cortical microstructure is gradually less mature at birth. Second, the observed age-related decreases in T_1_ and MD support the hypothesis of microstructural tissue growth in cortex during early infancy. A key question that remains is: what microstructural tissue compartments underlie this systematic postnatal cortical tissue growth?

### Gene expression data reveals myelination and synaptic processes are dominant mechanisms during early infancy

To answer this question, we leveraged the transcriptomic gene expression database of postmortem tissue samples of the Brain Span Atlas (https://www.brainspan.org) to identify candidate genes that show differential expression levels postnatally vs. prenatally. We reasoned that birth is a key developmental stage and genes that are expressed more in cortex postnatally than prenatally may contribute to the cortical development that we observed. We examined gene expression in brain tissue samples that closely matched our *in vivo* data in age (postnatal samples only) and anatomical location, including samples from primary sensory-motor cortices (M1, V1, A1, V1) and higher-level visual cortices (inferior parietal cortex, superior and inferior temporal cortex, **Supplementary Tables S6, S7**). For each sample, we extracted RNA-Seq expression data in Reads Per Kilobase per Million (RPKM) and determined which genes show significantly higher postnatal vs. prenatal expression along with the expression fold change (FC).

This differential analysis generated a list of several thousand genes that are expressed significantly more in these cortical expanses postnatally than prenatally. To determine the most differentially expressed genes, we selected the genes with the largest expression fold changes (FC > 4) and assessed their significance after Bonferroni correction for multiple comparisons (*P* < 5.7×10^−6^). **Fig. 3A** shows the expression level (per cortical sample/age) of ninety-five genes which survived these criteria and **Fig. 3B** shows their FC in descending order. For instance, for the top 10 differentially expressed genes, expression levels increase from ~1 RPKM, prenatally, to more than 6 RPKM, postnatally (**Fig. 3A**). Intriguingly, the top-most differentially expressed gene in primary sensory-motor and visual cortices is myelin basic protein (MBP), a gene associated with myelin generation and myelin sheath wrapping^22^. It is also interesting, that several other myelin-related genes, including myelin-associated oligodendrocytic basic protein (MOBP), myelin-associated glycoprotein (MAG), and proteolipid protein (PLP-1), are also among the top 10 most expressed genes postnatally (**Fig. 3A**).

**Fig 3.**
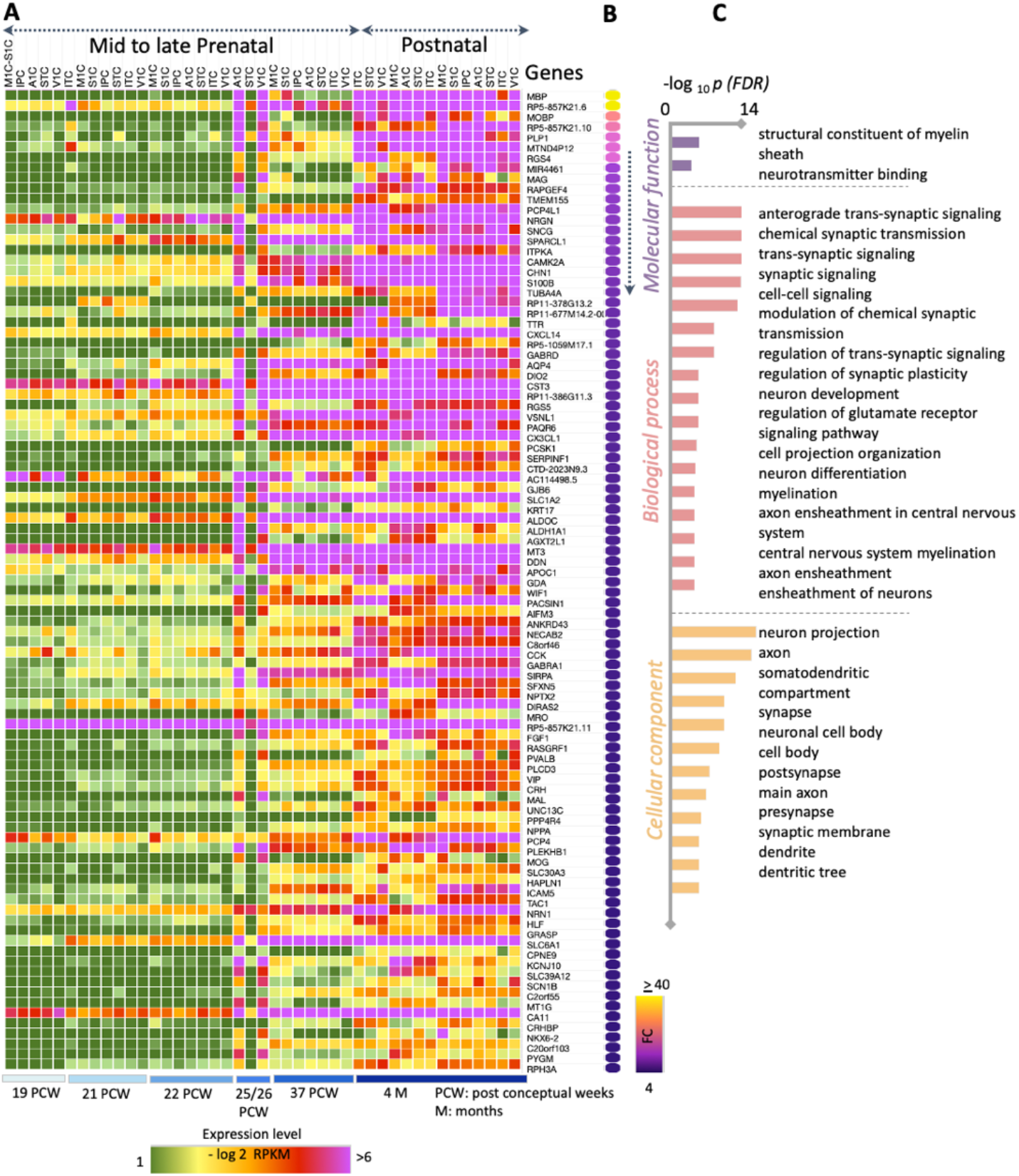
Transcriptomic gene analysis of cortical samples reveals that myelination and synaptic processes are cellular mechanisms of postnatal development. (A) Matrix showing gene expression levels in prenatal (19 post conceptual weeks (pcw) to 37 pcw) and postnatal (three 4-month-olds) cortical tissue samples for the 95 most differentially expressed genes. *Rows:* genes, *columns:* cortical area (acronyms in **Supplementary Table S7**), *color:* expression level in reads per kilobase per million (RPKM, see colorbar). (B) Gene expression fold change (FC) between postnatal vs. prenatal cortical samples of the 95 most differentially expressed genes. (C) Gene enrichment analysis^23^ showing the molecular and biological processes and cellular components related to these 95 genes.

To further elucidate the molecular and cellular pathways linked to these 95 genes, we used the ToppGene toolbox (https://toppgene.cchmc.org) to map this list of expressed genes to the enriched physiological processes. Comparing the 95 most significantly-expressed genes to all protein-coding genes as the background set, ToppGene reported significant enrichment of several biological processes related to: (i) myelination (P_FDR_corrected_=1.38×10^−4^), (ii) structural constituents of myelin sheath (P_FDR_corrected_=2.05×10^−5^), (iii) axonal ensheathing (P_FDR_corrected_=1.38×10^−4^), (iv) synaptic signaling (P_FDR_corrected_=1.34×10^−12^), and (v) cellular components of axonal projections and dendritic spines (P_FDR_corrected_=7.97×10^−5^) (**Fig. 3C, Supplementary Table S8** and **DataSet1**). These processes remained enriched in a control analysis in which we compared this list of top-95 most expressed genes to a different background gene set that was restricted to markers of cortical cells (neurons, astrocytes, endothelial cells, microglia, oligodendrocytes^24^ (**Supplementary DataSet2**).

### Early visual areas are more myelinated at birth, but myelinate at slower rates than later visual areas

Transcriptomic gene analyses indicated that myelination is a key microstructural component that is developing after birth. As higher myelin content decreases T_1_ relaxation time^14,15,17,18,25^ and mean diffusivity (MD) these data suggest that myelination contributes to the reduction in T_1_ and MD from birth to 6 months observed in cortex. This raises an intriguing possibility that these metrics might inform about the sequence and rate of cortical myelination in infancy. While MD depends on myelin, it also depends on axons’ radii and packing. As such, MD varies in complex nonlinear ways with myelin. Hence, it is difficult to make inferences from MD about cortical myelin^26^. T_1_ is related to myelin fraction^25^ 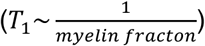. However, because of the inverse relationship, a change in T_1_ depends both on the change in myelin and the initial myelin fraction in the voxel^27^. Consequently, a similar change in myelin would produce a larger change in T_1_ in voxels that are less myelinated (smaller myelin fraction) than those that are more myelinated (larger myelin fraction). In contrast to T_1_, tissue relaxation rate 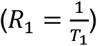 varies linearly with myelin^25^. Therefore, changes in R_1_ are linearly related to myelin changes, independent of a voxel’s myelin fraction. Hence, to glean insights about cortical myelin in early infancy, we measured R_1_ in each infant, timepoint, and visual area.

Results revealed linear increases in R_1_ in both ventral and dorsal visual processing streams and in both hemispheres (**Supplementary Fig. S8**). We used LMMs to evaluate R_1_ at birth and change in R_1_ per day across the first 6 months of life. We found significant differences across visual areas of R_1_ at birth (**Fig. 4A,C**). Using a random intercept/random slope LMM relating the mean R_1_ to participants’ age [in days] across all ROIs of a stream (see Methods), we found a significant effect of ROI across both streams and hemispheres (*ts* > 5.00, *Ps* < 1.102×10^−8^). Like T_1_ at birth (**Fig. 2**), which varied progressively across the visual hierarchy, R_1_ at birth in the right hemisphere is highest in V1 (R_1_=0.49±0.007) and progressively decreased from early to later visual areas across both visual streams (**Fig. 4A,C)**. Indeed, R_1_ at birth is numerically lowest in right IPS1 (R_1_=0.42±0.005) in the dorsal stream and right V02 (R_1_=0.44±.005) in the ventral stream (**Supplementary Table S9**). Surprisingly, the change in R_1_ shows an opposite pattern, being progressively larger in later than earlier visual areas of both visual streams (**Fig. 4B,D, Supplementary Table S9**). E.g., change in R_1_ is 0.59 ± 0.06 [ms^−1^/day] in right dorsal V1d versus 0.76 ± 0.04 [ms^−1^/day] in right IPS0 (first visual region of the intra parietal sulcus). In other words, R_1_ in IPS0 develops ~28% faster than in V1 during the first 6 months of life. We observed a similar pattern in the ventral stream: R_1_ changes at a rate of 0.55 ± 0.06 [ms^−1^/day] in V1v and at rate of 0.76 ± 0.04 [ms^−1^/day] in VO2. As R_1_ is linearly related to myelin, and cortical iron is negligible in early infancy^28^, our data suggest that in both processing streams, earlier visual areas have higher myelin content at birth, but they myelinate at slower rates than later visual areas.

**Fig 4.**
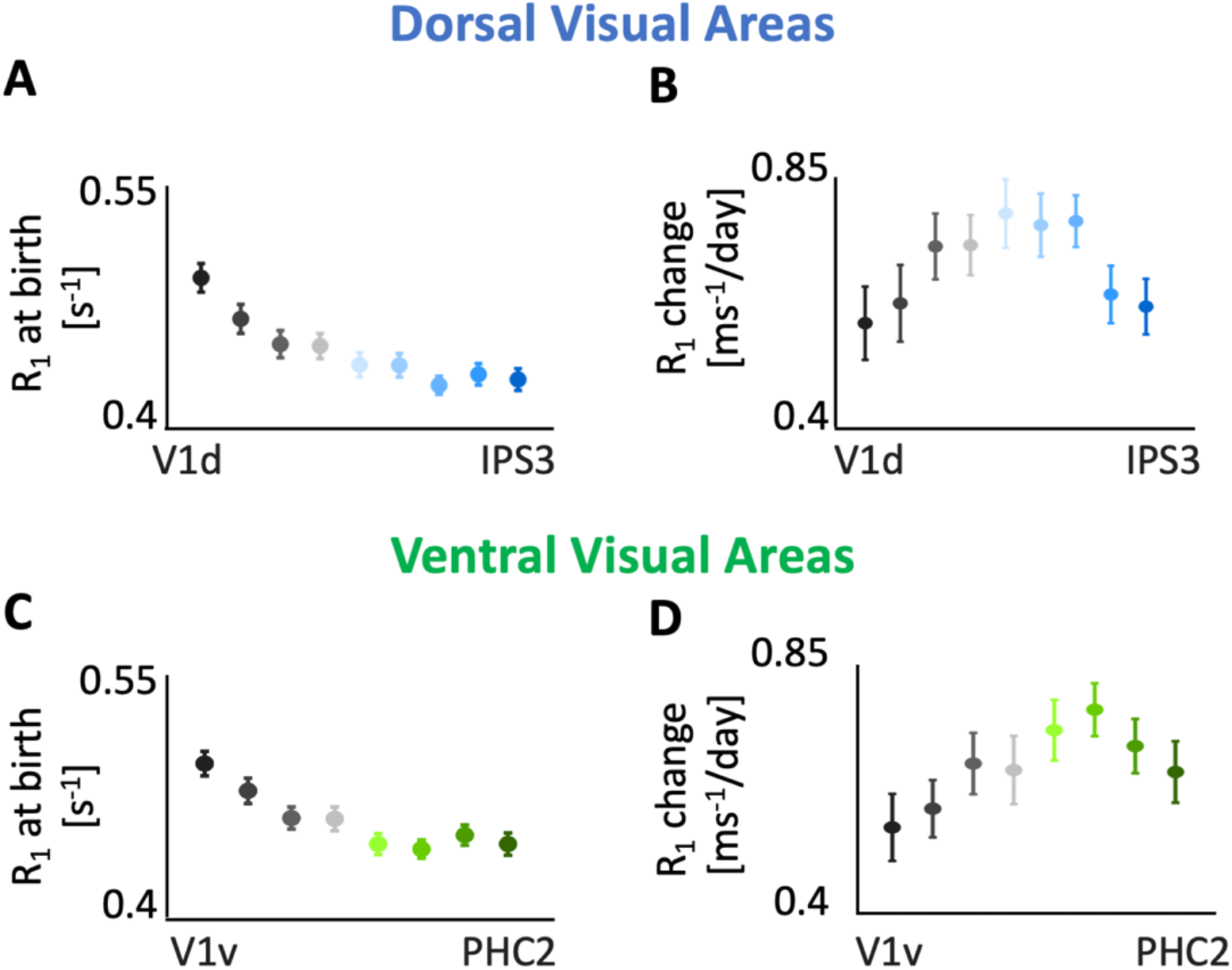
Both R_1_ at birth and change in R_1_ per day vary systematically across visual processing streams. Linear mixed modeling (LMM) of R_1_ as a function of age (**Supplementary Fig. S8)** estimate R_1_ at birth (LMM intercept) and rate of R_1_ development (LMM slope) for each visual area in the dorsal (A,B) and ventral (C,D) visual streams. *Error bars:* standard error on estimates of intercepts and slopes. Data show are from right hemisphere; Left hemisphere data in **Supplementary Fig. S8**.

## Discussion

Combining innovative *in vivo* metrics that have meaningful units with gene expression analyses, we show that during the first 6 months of life, infants’ cortex undergoes exuberant microstructural tissue growth related to myelination, synaptogenesis, and dendritic processes. Within visual cortex, we find hierarchical development across two processing streams, where earlier regions are more mature at birth, but develop slower than later visual areas.

We find no evidence for cortical tissue pruning in sensory-motor cortices from 0 to 6 months of age. This is consistent with *ex vivo* data showing synaptogenesis^2^ and dendritic growth^10^ in primate V1 and A1 during early infancy. Nonetheless, it is possible that pruning happens later in development after this exuberant growth^2,29^, or that pruning effects are smaller than microstructural growth effects, which dominate MRI metrics.

Gene analyses reveal that MBP is the top-most differentially expressed gene after birth, and both *ex vivo* and *in vivo* studies show that cortical R_1_ increases with higher myelin content^15,18^. Consequently, this suggests that myelination contributes to tissue changes in infants’ cortex. These findings are transformative for two reasons. First, they link specific biological mechanisms to *in vivo* MRI measurements. Second, they lead to a new hypothesis that cortical myelination is critical for the development of brain function and ultimately behavior. Thus, we believe that any infant research^30–32^, should consider the impact of cortical myelination on development, which has largely been overlooked.

Finally, our results overturn the prevailing theory that primary visual cortex is myelinated at birth^1,33^ and develops faster than higher-level areas^2^. Our analyses of R_1_ not only suggest that both early and later visual areas continue to develop and myelinate during the first six months of life, but also that contrary to prevailing hypothesis early sensory-motor areas do not myelinate the fastest. We hypothesize that more mature primary sensory-motor cortices at birth provide scaffolding for the development of cortical systems, but the sensory richness of the postnatal environment accelerates activity-dependent myelination and synaptogenesis in higher-level sensory cortices^9,34,35^.

In conclusion, our findings necessitate a rethinking of how cortical microstructure develops in infants, and open new avenues to examine the impact of cortical myelination on the development of brain function. Young infants are a highly vulnerable population for which the ability to diagnose delayed and atypical development could not be of greater importance. These novel, powerful multimodal methodologies enable non-invasive, longitudinal measurements within individual infants that are linked to specific biological mechanisms. Thus, our study has important implications for identifying neurodevelopmental delays and disorders in infants which may lead to early inventions and better life-long outcomes.

## Methods

### Participants

16 full-term and healthy infants (7 female) were recruited to participate in this study. 3 infants provided no usable data because they could not stay asleep once the MRI sequences started. Here, we report data from 13 infants (6 female) across three age timepoints: newborn/0 months [8-37 days], 3 months [78-106 days], and 6 months [167-195 days], with 10 participants per timepoint. Two participants were reinvited to complete scans for their 6-months session that could not be completed during the first try. Both rescans were performed within 7 days and participants were still within age range for the 6-month timepoint. The participant population was racially and ethnically diverse reflecting the population of the San Francisco Bay Area, including two Hispanic, nine Caucasian, two Asian, three multiracial (2 Asian and Caucasian; 1 Native Hawaiian or Other Pacific Islander) participants. Seven out of these 13 infants participated in MRI in all three timepoints (0, 3, 6 months). Due to the Covid-19 pandemic and restricted research guidelines, data acquisition was halted. Consequently, the remaining infants participated in either 1 or 2 sessions. Participation of the 13 infants whose data is reported in this study is summarized in **Supplementary Table S1**.

#### Expectant mother and infant screening procedure

Expectant mothers and their infants in our study were recruited from the San Francisco Bay Area using social media platforms. We performed a two-step screening process for expectant mothers. First, mothers were screened over the phone for eligibility based on exclusionary criteria designed to recruit a sample of typically developing infants and second, eligible expectant mothers were screened once again after giving birth. Exclusionary criteria for expectant mothers were as follows: recreational drug use during pregnancy, significant alcohol use during pregnancy (more than 3 instances of alcohol consumption per trimester; more than 1 drink per occasion), lifetime diagnosis of autism spectrum disorder or a disorder involving psychosis or mania, taking prescription medications for any of these disorders during pregnancy, insufficient written or spoken English ability to comprehend study instructions, and learning disabilities. Exclusionary criteria for infants were preterm birth (<37 gestational weeks), low birthweight (<5 lbs. 8 oz), small height (<18 inches), any congenital, genetic, or neurological disorders, visual problems, complications during birth that involved the infant (e.g., NICU stay), history of head trauma, and contraindications for MRI (e.g., metal implants).

### MRI (magnetic resonance imaging) Procedure

We acquired T2-weighted MRI, quantitative MRI (qMRI), and diffusion MRI (qMRI) data for each infant and time point. Study protocols for these scans were approved by the Stanford University Internal Review Board on Human Subjects Research. Scanning sessions were scheduled in the evenings close in time to the infants’ typical bedtime. The duration of each session lasted between 2.5 – 5 hours including time to prepare the infant and waiting time for them to fall asleep. Upon arrival, caregivers provided written, informed consent for themselves and their infant to participate in the study. Before entering the MRI suite, both caregiver and infant were checked to ensure that they were safe and metal-free to enter the MRI suite. Before entering the MRI suite, caregivers changed the infants into MR safe cotton onesies and footed pants provided by the researchers. The infant was swaddled with a blanket with their hands to their sides to avoid their hands creating a loop during MRI. During scans of newborn infants, an MR safe plastic immobilizer (MedVac, www.supertechx-ray.com) was used to stabilize the infant and their head position. Once the infant was ready for scanning, the caregiver and infant entered the MR suite. The caregiver was instructed to follow their child’s typical sleep routine. As the infant was falling asleep, researchers inserted soft wax earplugs into the infant’s ears. Once the infant was asleep, the caregiver was instructed to gently place the infant on a makeshift cradle on the scanner bed, created by placing weighted bags at the edges of the bed to prevent any side-to-side movement. Finally, to lower sound transmission, MRI compatible neonatal Noise Attenuators (https://newborncare.natus.com/products-services/newborn-care-products/nursery-essentials/minimuffs-neonatal-noise-attenuators) were placed on the infant’s ears and additional pads were also placed around the infant’s head to stabilize head motion. An experimenter stayed inside the MR suite with the infant during the entire scan in case the infant woke up during scanning. For additional monitoring of the infant’s safety and motion quality an infrared camera was affixed to the head coil and positioned for viewing the infant’s face in the scanner. The researcher operating the scanner monitored the infant via the camera feed, which allowed for the scan to be stopped immediately if the infant showed signs of waking or distress. This setup allowed tracking the infant’s motion; scans were stopped and repeated if there was excessive head motion.

#### Data Quality Assurance during MRI

To ensure high data quality, in addition to real-time monitoring of the infant’s motion via an infrared camera, acquired scans were assessed immediately after acquisition of each sequence and repeated if necessary. Factors for repetition included head motion detected on the infrared camera, or head motion detected on the acquired MR images reflected in blurring of otherwise detailed anatomical images and partial voluming effects. On average, 50% of all scans across infants were successful on the first try while the rest had to be repeated.

### Data Acquisition

All participants participated in multiple scans in each session to obtain anatomical MRI, quantitative MRI (qMRI), and diffusion MRI (dMRI) data. Data were acquired at two identical 3T GE Discovery MR750 Scanners (GE Healthcare) and Nova 32-ch head coils (Nova Medical) located at Stanford University: (i) Center for Cognitive and Neurobiological Imaging (CNI) and (ii) Lucas Imaging Center. As infants have low weight, all imaging was done with Normal SAR level to ensure their safety.

#### Anatomical MRI

##### T2-weighted

T2-weighted images were acquired for participants in each of 0, 3, 6 months timepoints. T2-weighed image acquisition parameters: TE=124 ms; TR = 3650ms; echo train length = 120; voxel size = 0.8mm^3^; FOV=20.5cm; Scan time: 4 min and 5 sec.

#### Quantitative MRI

##### Spoiled-gradient echo images (SPGR)

were acquired for all participants in each of 0, 3, 6 months. These images were used together with the IR-EPI sequence to generate whole-brain synthetic T1-weighted images. We acquired 4 SPGRs whole brain images with different flip angles: α = 4°, 10°, 15°, 20°; TE=3ms; TR =14ms; voxel size=1mm^3^; number of slices=120; FOV=22.4cm; Scan time: (4:55 min) x 4.

#### Inversion-recovery EPI (IR-EPI)

IR-EPI images were acquired from all participants in each of 0, 3, 6 months timepoints. We acquired multiple inversion times (TI) in the IR-EPI using a slice-shuffling technique^36^: 20 TIs with the first TI=50ms and TI interval=150ms; we also acquired a second IR-EPI with reverse phase encoding direction. Other acquisition parameters are voxel size = 2mm^3^; number of slices=60; FOV=20cm; in-plane/through-plane acceleration = 1/3; Scan time=1:45 min) x 2.

#### Diffusion MRI

We obtained dMRI data from 9 newborn participants, and 10 participants at 3 months and 6 months of age. One newborn woke up prior to run completion and we could not complete dMRI acquisition. dMRI parameters: multi-shell, #diffusion directions/b-value = 9/0, 30/700, 64/2000; TE = 75.7 ms; TR=2800ms; voxel size = 2mm^3^; number of slices=60; FOV=20cm; in-plane/through-plane acceleration = 1/3; Scan time: 5:08 min. We also acquired a short dMRI scan with reverse phase encoding direction and only 6 b=0 images (scan time 0:20 min).

### Data Analysis

The data analysis pipeline is summarized in **Supplementary Fig. S1**. In brief, IR-EPI data were used to estimate T_1_ relaxation time at each voxel. These data were also used together with the SPGRs to generate synthetic T1-weighted whole brain anatomies of each infant at each timepoint. All data from that timepoint were aligned to this anatomical image. T2-weighted images were used for segmentation of gray-white matter to generate cortical surface reconstructions and dMRI data were used to estimate MD in each voxel. All infant data were kept in native space as all analyses were performed within-subject and within-timepoint.

#### Quantitative T_1_ relaxation time modeling

The signal equation of *T*_*1*_ relaxation of an inversion-recovery sequence is an exponential decay:

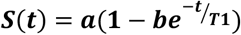

where *t* is the inversion time, *a* is proportional to the initial magnetization of the voxel, *b* is the effective inversion coefficient of the voxel (for perfect inversion *b=2*). To work with magnitude images, we took the absolute value of the above signal equation and used it as the fitting model.

First, as part of the preprocessing, we performed susceptibility-induced distortion correction on the IR-EPI images using FSL’s^37^ top-up correction and the IR-EPI acquisition with reverse phase encoding direction. We then used the distortion corrected images to fit the above T_1_ relaxation signal model using a multi-dimensional Levenberg-Marquardt algorithm^38^. The output of the algorithm is the estimated T_1_ in each voxel as well as the model goodness of fit (R^2^) value in each voxel.

#### Generation of T1-weighted whole brain anatomies

From the SPGRs and IR-EPI scans, synthetic T1-weighted whole brain images were generated using mrQ software (https://github.com/mezera/mrQ). We analyzed all data in the native infant space and did not align to any template brain. All data (T2-weighted-anatomy, quantitative T_1_ relaxation estimates, and MRI data) from a given timepoint were aligned to this brain volume (**Supplementary Fig. S1**).

#### Generation of cortical surfaces

To generate cortical surface reconstructions, we used both T2-weighted and synthetic T1-weighted anatomies. We used multiple steps to generate accurate cortical surface reconstructions of each infant’s brain at each timepoint. (1) An initial segmentation of gray and white matter was generated from the synthetic T1-weighted brain volume using infant FreeSurfer’s automatic segmentation code developed for infant data (infant-recon-all; https://surfer.nmr.mgh.harvard.edu/fswiki/infantFS)^39^. This initial segmentation generates pial and white matter surfaces, and surfaces of curvature, thickness, surface area. However, this initial FreeSurfer’s segmentation misses significant portions of the infant’s gray matter, as the contrast of infants’ T1-weighted images were not differentiated enough between gray and white matter to generate an accurate segmentation. (2) We used T2-weighted anatomical images, which have a better contrast between gray and white matter in infants, and an independent brain extraction toolbox (Brain Extraction and Analysis Toolbox, iBEAT, v:2.0 cloud processing, https://ibeat.wildapricot.org/) to generate more accurate white and gray matter segmentations. (3) The iBEAT segmentation was further manually corrected to fix segmentation any additional errors (such as holes and handles) using ITK-SNAP (http://www.itksnap.org/) in white matter as well as gray matter. (4) The manually corrected iBEAT segmentation was aligned to the T1-weighted anatomy that was used for the FreeSurfer segmentation using manual rigid body alignment in ITK-SNAP (**Supplementary Fig. S1**). (5) This aligned and segmented volume was then reinstalled into FreeSurfer using software we developed based on infant FreeSurfer functions (https://github.com/VPNL/babies_graymatter). This process updates the white matter segmentation and the cortical surfaces in the subject’s FreeSurfer directory to render the accurate surfaces (**Supplementary Fig. S1**). This accurate surface was used for visualization, and cortex-based registration with atlases.

#### dMRI

DMRI data were preprocessed using a combination of tools from mrTrix3 (https://github.com/MRtrix3/mrtrix3)^40^ and mrDiffusion toolbox (http://github.com/vistalab/vistasoft). (1) We denoised the data using a principal component analysis^41^ (2) We used FSL’s top-up tool (https://fsl.fmrib.ox.ac.uk/) and one image collected in the opposite phase-encoding direction to correct for susceptibility-induced distortions. (3) We used FSL’s eddy to perform eddy current and motion corrections. Motion correction included outlier slice detection and replacement^42^ (4) We performed bias correction using ANTs^43^ (5) These preprocessed dMRI images were registered to the whole-brain T2-weighted anatomy using whole-brain rigid-body registration in a two-stage model with a coarse-to-fine approach that maximized mutual information. (6) mrTrix3 software was used to fit tensors to each voxel using a least-squares algorithm that removes outliers. From the kurtosis tensor files, we estimated mean diffusivity (MD) maps in each voxel of the brain.

#### dMRI quality assurance

Out of the 29 dMRI acquisitions, one newborn acquisition was missing the reverse phase encoding image required for susceptibility correction. Thus, we report data from eight newborns, ten 3-month-olds and ten 6-month-olds. Across all acquisitions, less than 5.0% ± 0.7% of dMRI images were identified as outliers by FSL’s eddy tool. We found no significant effect of age across the outliers (no main effect of age: F_2,25_=2.84, *P*=0.08, newborn:1.20±0.83; 3 months:0.4±0.40; 6 months: 0.67±0.85) suggesting that the developmental data was well controlled across all ages of infants.

### Delineation of the primary sensory cortices and ventral and dorsal visual areas

Our goal was to examine developmental changes in quantitative T_1_ and MD in the gray matter of four primary sensory-motor cortices as well as across the ventral and dorsal visual processing streams from 0 to 6 months of age. In order to delineate these regions in infants, we used brain atlases that delineate these regions. Each atlas is available on the FreeSurfer adult average brain and was projected to each individual participant’s cortical surface at each timepoint using infant FreeSurfer’s cortical-based alignment tool. Without functional data, cortex-based alignment is the most accurate method for defining brain areas in individual brains from atlases^44^. Critically, the major sulci and gyri that are used for cortex-based alignment are present at birth^45^ and are visible on the cortical surface reconstructions of each of our infants. We used the Glasser atlas^21^ to delineate the primary visual (V1), primary auditory (A1), primary motor (M1), and primary somatosensory (S1) cortices. We used the Wang atlas^20^ to delineate 9 regions spanning the dorsal visual stream (V1d, V2d, V3d, V3a, V3b, IPS0, IPS1, IPS2, and IPS3) and 8 regions spanning the ventral visual stream (V1v, V2v, V3v, hV4, V01, V02, PHC1, and PHC2) in each participant’s brain.

#### Quality check on the delineation of regions of interest (ROIs)

As we used cortex-based alignment and atlases developed for the adult brain to define areas in the infant brain, we sought to test how well the ROIs identified in infants’ brains compared to those identified in adults’ brains. We reasoned that if regions identified from adult atlases would show better correspondence to manually defined regions in adults than infants, it would indicate that these atlases are in optimal for infants. To test this, we compared how the calcarine sulcus manually defined on each individual brain compares to the cortex based aligned calcarine sulcus from the Desikan atlas (an anatomical parcellation of the brain, in the average adult FreeSurfer cortical surface^46^. The manually defined calcarine sulcus was defined from the posterior end of the occipital pole to the prostriate (ProS) - a region along the anterior bank of the parietal-occipital sulcus. We chose the calcarine sulcus as the benchmark because V1 is located in the calcarine sulcus and it can be identified anatomically in each individual brain. In each infant and timepoint (*N*=30) and 10 adults (ages 22-27; from our previous study^18^), we calculated the overlap between the Desikan-calcarine and individual-subject-calcarine using the dice coefficient^47^. We found that the dice coefficient was 0.66±0.10 (Mean±SD) in infants and 0.67±.05 in adults. There were no significant age-related differences in dice-coefficients between infants and adults (no significant main effect of age: F_1,76_ =.02, *P*=.89, 2-way analysis of variance (ANOVA) with factors of age group (infant/adult) and hemisphere (left/right) and no significant differences between hemispheres (F_1,76_ =.02, *P*=.88). This analysis suggests that cortex-based alignment of brain atlases based on adult templates to infants’ cortical surfaces is similar in quality to this transformation in adults.

### Analysis of mean T_1_, MD, and R_1_ in cortical areas and their development

After delineating brain areas in each participant, we calculated the distribution and mean T_1_ / R_1_ and mean diffusivity (MD) in each participant and timepoint. We used linear mixed models (LMMs) to determine if there were age-related changes of T_1_ and MD within and across areas. LMMs were fit using the MATLAB 2017b function *fitlme*. In order to quantify the development of T_1_ / R_1_ and MD, for each area we fit a LMM relating the mean metric of that area to participants’ age [in days]. We ran two types of LMM per area and data type: (1) LMM with random intercept/fixed slope model, which allows only the intercepts to vary across participants and accounts for the fact that the same infants participated across multiple timepoints, and (2) LMM with random intercept/random slope model, which allows both intercepts and slopes to vary across participants. Results revealed that the random intercept/fixed slope model fitted the data best in all cases (in all model comparisons: Ps<0.05). Thus, we report the parameters of the LMMs with random intercepts in all our analyses below:

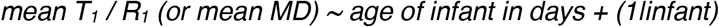

where *Mean T*_*1*_ */ R*_*1*_ *(MD)* is the dependent variable, age is a continuous predictor (fixed effect), and the term: 1 | infant indicates that random intercepts are used for each participant. Per model we obtained: (1) the intercept, which represented the values of *T*_*1*_ *(MD)* at birth, and (2) the average slope, which represented the rate of *T*_*1*_ *(MD*) development. The linear fits of the LMMs are plotted in **Figs. 1–2** and **Supplementary Figs. S3-S6** and we report slopes, and significant levels (Ps) of all areas in both hemispheres in **Supplementary Tables S2-S5** below. Since we ran LMMs for each area individually, we performed Bonferroni correction for multiple comparisons for each analysis: (1) across primary sensory-motor cortices (4 areas, 6 comparisons), (2) across the dorsal visual stream (9 areas, 36 comparisons), and (3) across the ventral visual stream (8 areas, 28 comparisons).

We used a second set of the linear mixed models to quantify the development in T_1_, R_1,_ MD across all ROIs within a stream. We fit a LMM relating the mean metric (T_1_ or R_1_ or MD) of that area to participants’ age [in days]. We ran two types of LMM per stream and metric: (1) LMM with random intercept/fixed slope model, which allows only the intercepts to vary across ROIs, and (2) LMM with random intercept/random slope model, which allows both intercepts and slopes to vary across ROIs. Results revealed that the random intercept/random slope model fitted the data best for T_1_ and R_1_ (in all model comparisons the latter model was significantly better than the former, likelihood test, Ps<0.01, except for trend in the left ventral stream, Ps>.15; degrees of freedom (ventral): 236; degrees of freedom (dorsal): 266). Thus, we report the parameters of the LMMs with random intercepts/slopes for these metrics. For MD, results revealed that the random intercept/fixed slope model fitted the data best (in all model comparisons the latter model was not significantly better than the former, Ps>0.05, except for a significant difference for the left dorsal stream, likelihood test, P=0.01, degrees of freedom (ventral): 220; degrees of freedom (dorsal): 248). Thus, we report the parameters of the LMMs with random intercepts/fixed slopes for MD.

### Transcriptomic gene data analysis of postnatal versus prenatal tissue samples

In order to assess what microstructural tissue compartments may be linked to the observed postnatal tissue growth in cortex related to decrease in T_1_ and MD, we used the transcriptomic gene expression database of post-mortem tissue samples made available by the Brain Span Atlas (https://www.brainspan.org). We examined: (1) if there were differences in the expression levels for genes in the postnatal tissue samples as compared to the prenatal tissue samples and (2) if so, what cellular and biological processes are related to these genes of interest. In order to closely match our *in vivo* data, we selected the postmortem postnatal tissue samples within our *in vivo* age range (0 to 6 months). Prenatal samples were between 19 post conceptual weeks (pcw) to 37 pcw, which is just prior to birth. **Supplementary Table S6** includes demographic details of the postnatal and prenatal samples. Within these postmortem samples, we compared tissue from primary sensory cortices (V1, M1, S1, and A1) and parietal and temporal regions overlapping visual regions of the dorsal and ventral visual streams, respectively, to match our *in vivo* data (**Supplementary Table S7** for the complete list of brain regions from individual prenatal and postnatal samples).

The differential analysis provides information about which genes are differentially expressed when we compared our target (postnatal) versus control (prenatal) sample sets. Specifically, differential expression-level analysis reveals a list of several thousand genes along with the gene level RNA-Sequencing expression data in reads per kilobase per million (RPKM, data were log2-scaled). The analysis also estimated how many more times the genes were expressed postnatally vs prenatally (fold changes, FC) and provides the statistical significance of the contrast (*p-*values). Fold change is measured as the average log2(intensity/expression) values of all samples in the target sample minus the average log2(intensity/expression) of the control samples. As standard practice^48^ we applied two thresholds: (1) a threshold of fold change: FC>4) and (2) a Bonferroni correction (P<5.7×10^−6^) related to the differential analysis. Ninety-five genes passed these thresholds.

#### Functional enrichment

Next, to assess what molecular and biological processes are linked to our top gene list we inputted the list of genes to a toolbox created for gene list enrichment analysis (ToppGene https://toppgene.cchmc.org) 23 Specifically, this toolbox identifies the biological pathways that are enriched (over-represented) by the expression of the genes of interest more than that would be expected by chance. Each gene is compared with the genes related to a specific pathway and a p-value of the enrichment of a pathway is computed and multiple-test correction is applied. A table of biological, molecular, and cellular processes related to this list of genes is derived from this analysis. A Benjamini–Hochberg False Discovery Rate (FDR) was applied as a multiple comparisons adjustment. Information on the biological pathways related to the top-most genes, is ranked by statistical significance of functional enrichment (−log_10_ (p-value FDR). We performed this analysis first using (i) all protein-coding genes in the ToppGene database as the background reference set and (ii) only including genetic markers of cortical cells as the background reference set (N_genes_=5000) including neurons, astrocytes, endothelial cells, microglia, oligodendrocytes^24^ see **Supplementary Table S8** for the FDR corrected p-values of the molecular functions, biological processes, and cellular processes listed in **Fig. 3c**. Complete gene ontology lists without and with background gene sets can be found on GitHub: https://github.com/VPNL/babies_graymatter/genes/Dataset1 and https://github.com/VPNL/babies_graymatter/genes/Dataset2.

## Acknowledgements

We would like to acknowledge Amy Kang, KK Barrows, Javier Marquis Lopez, Laura Villalobos, Lois Williams, and Alex Rezai for their contributions with gray and white matter segmentations of the infant brains and Caitlyn Estrada for her contribution to data collection. We would also like to thank Jiyeong Ha for her contributions towards the quality assurance check on the delineations of brain regions on the infant and adult brains. Additionally, we would like to thank Fiorella Carla Grandi for feedback and advice on the gene analyses.

## Author contributions

VSN: data preprocessing, statistical analysis, manuscript writing; MR: study design, data acquisition, data preprocessing, statistical analysis, manuscript writing; HW: sequence development; FRQ: participant recruitment, data acquisition, data preprocessing; HK: participant recruitment, data acquisition, data preprocessing;; NLA: data collection; MG: data preprocessing (dMRI), SB: data preprocessing (artificial T1-weighted image), AAM: sequence development, KGS: designed the study, oversaw all components of the study, and wrote the manuscript. All co-authors read and approved the submitted manuscript.

## Data and materials availability

Data, code, and materials used in the analysis will be made available with publication on GitHub (https://github.com/VPNL/babies_graymatter).

---

**Supplementary Figure 1.**
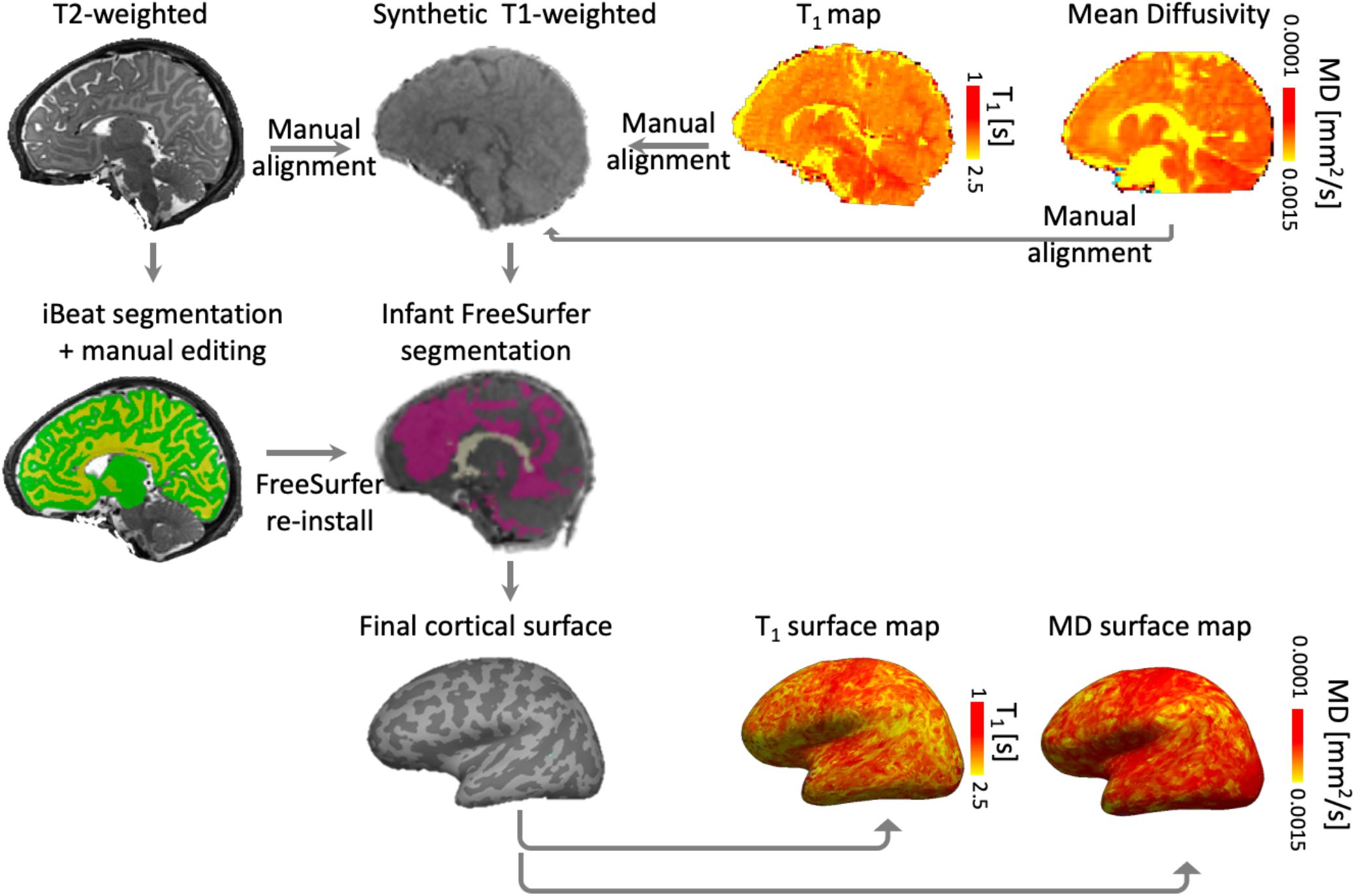
MRI data preprocessing pipeline. Schematic showing the preprocessing pipeline associated with obtaining the white and gray matter segmentations for generating the cortical surfaces and quantitative T_1_ and mean diffusivity (MD) maps per individual baby brain.

**Supplementary Figure 2.**
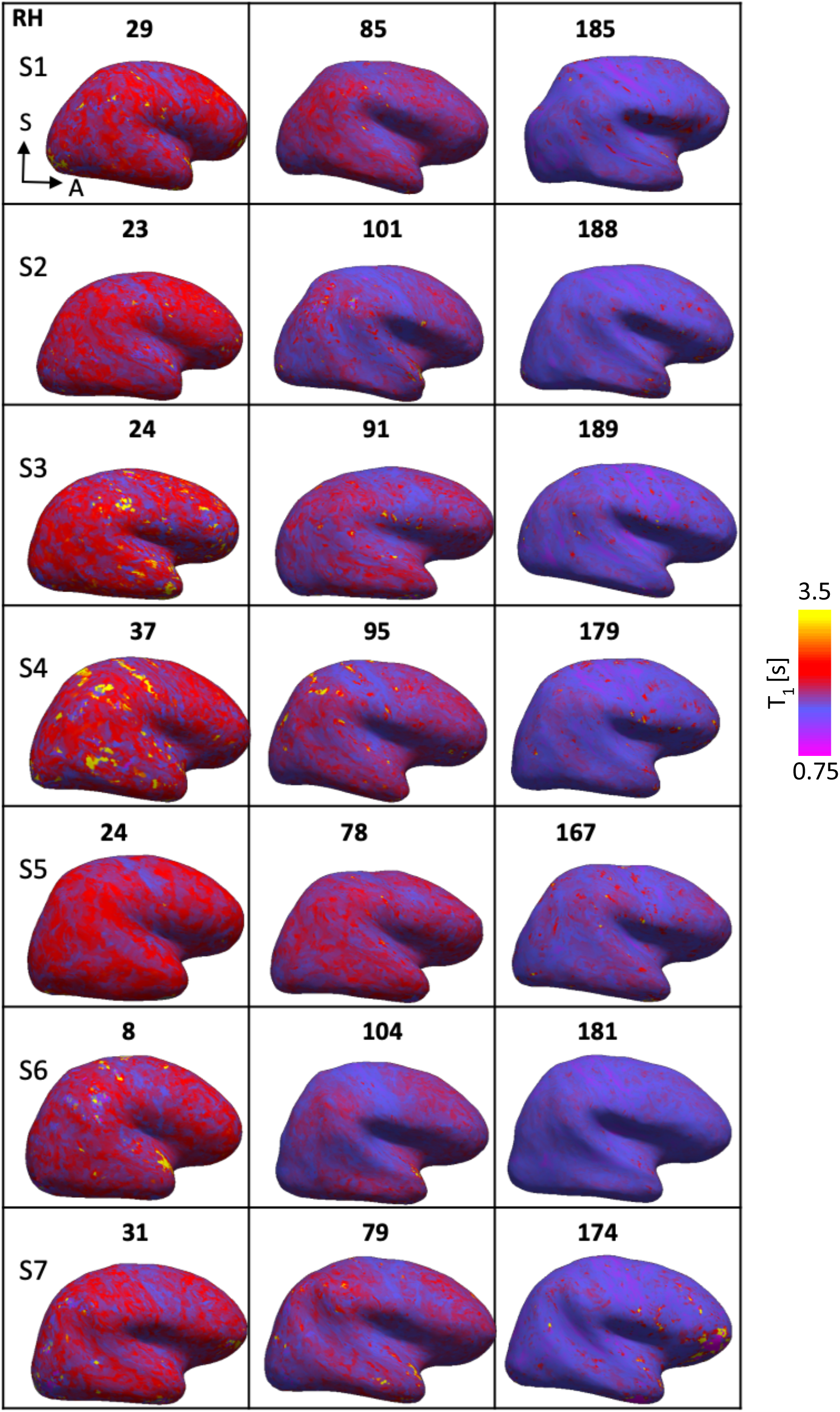
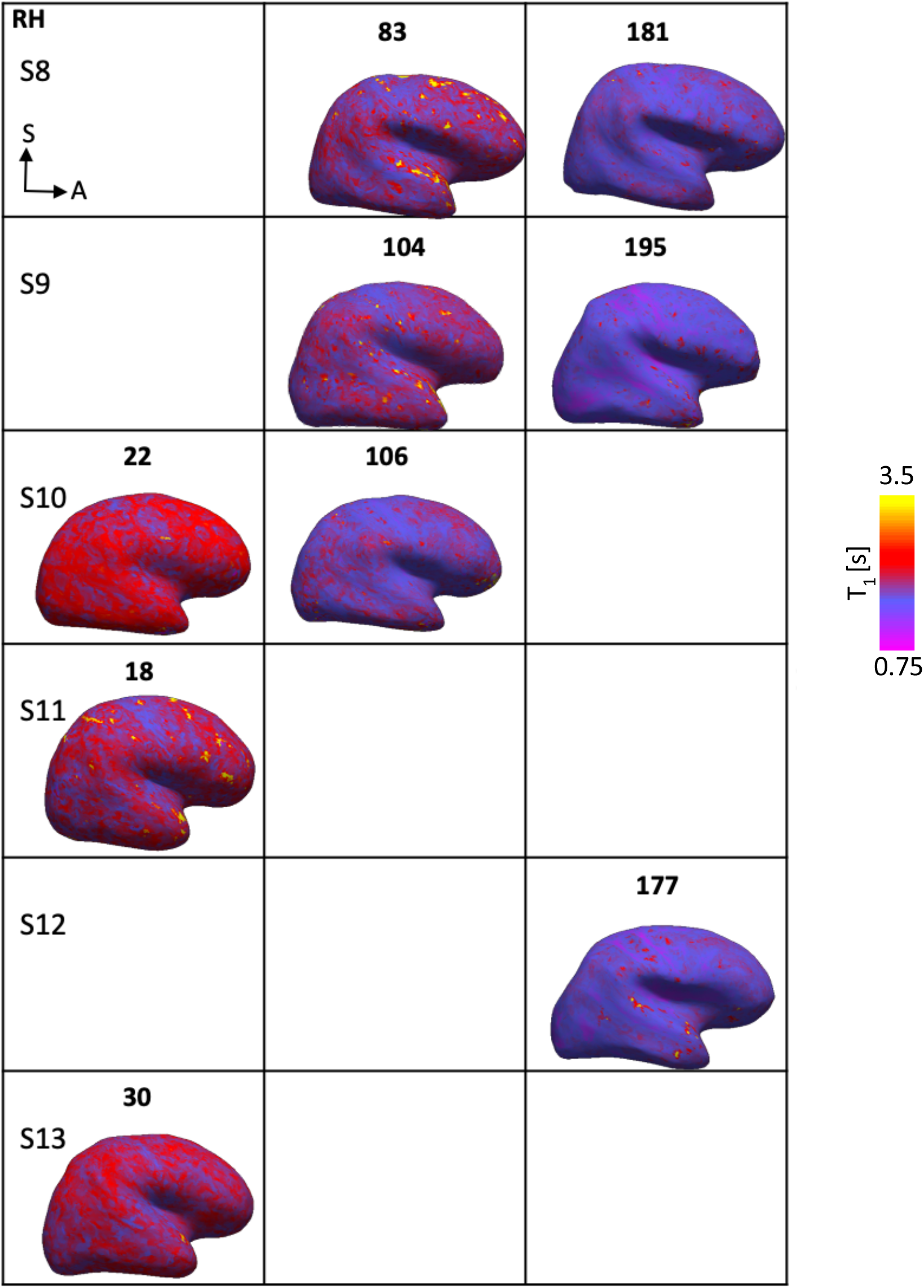
Right cortical surfaces in the lateral view showing T_1_ development in seven individual babies across three time points of development: newborns, and approximately 3 and 6 months of age. Each row represents an individual baby. T_1_ [s] decreases from newborn (red) to 6 months (purple) of age. Age at scan (in days) isindicated above each cortical surface. *RH: right hemisphere, S: superior, A: anterior*.

**Supplementary Figure 3.**
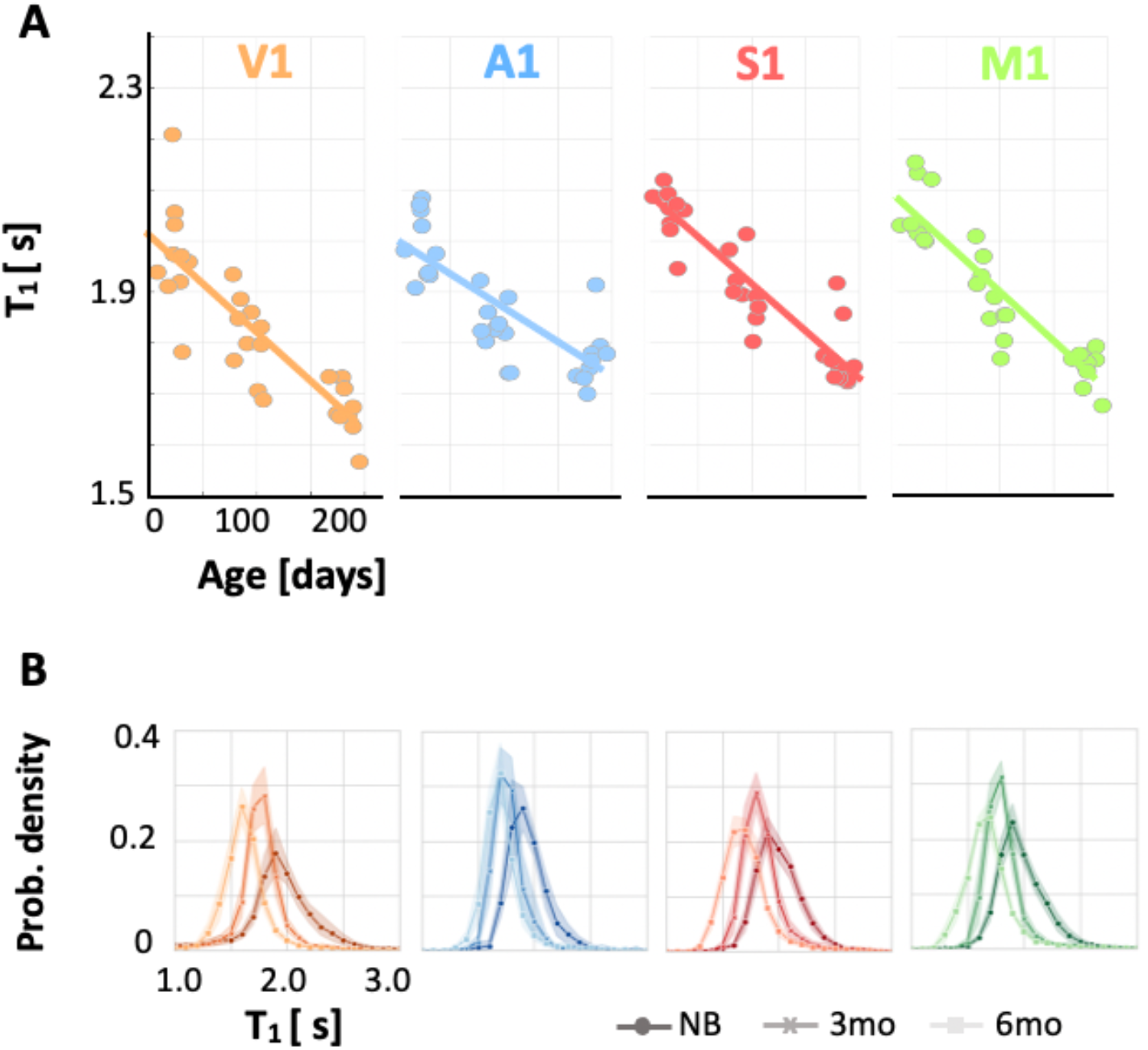
Mean T_1_ in four early sensory regions in the left hemisphere decreases with age in the first six months of infant life. (A) Significant decrease in T_1_ with age in the four primary sensory-motor cortices (*V1: primary visual cortex, A1: primary auditory cortex, S1: primary somatosensory cortex, M1: primary motor cortex*); slopes and significance (p-values) in **Supplementary Table S2**. Each dot represents mean T_1_ of an infant’s region of interest. (B) Distribution of T_1_ across voxels in each area decreases from newborns (darker colors) to 6-month-olds (lighter colors). *Solid lines:* mean distribution across participants; *shaded region:* standard error of the mean (SE) across 10 infants in each timepoint. *NB:* newborns; *3 mo:* 3 month-olds; *6 mo:* 6 month-olds.

**Supplementary Figure 4.**
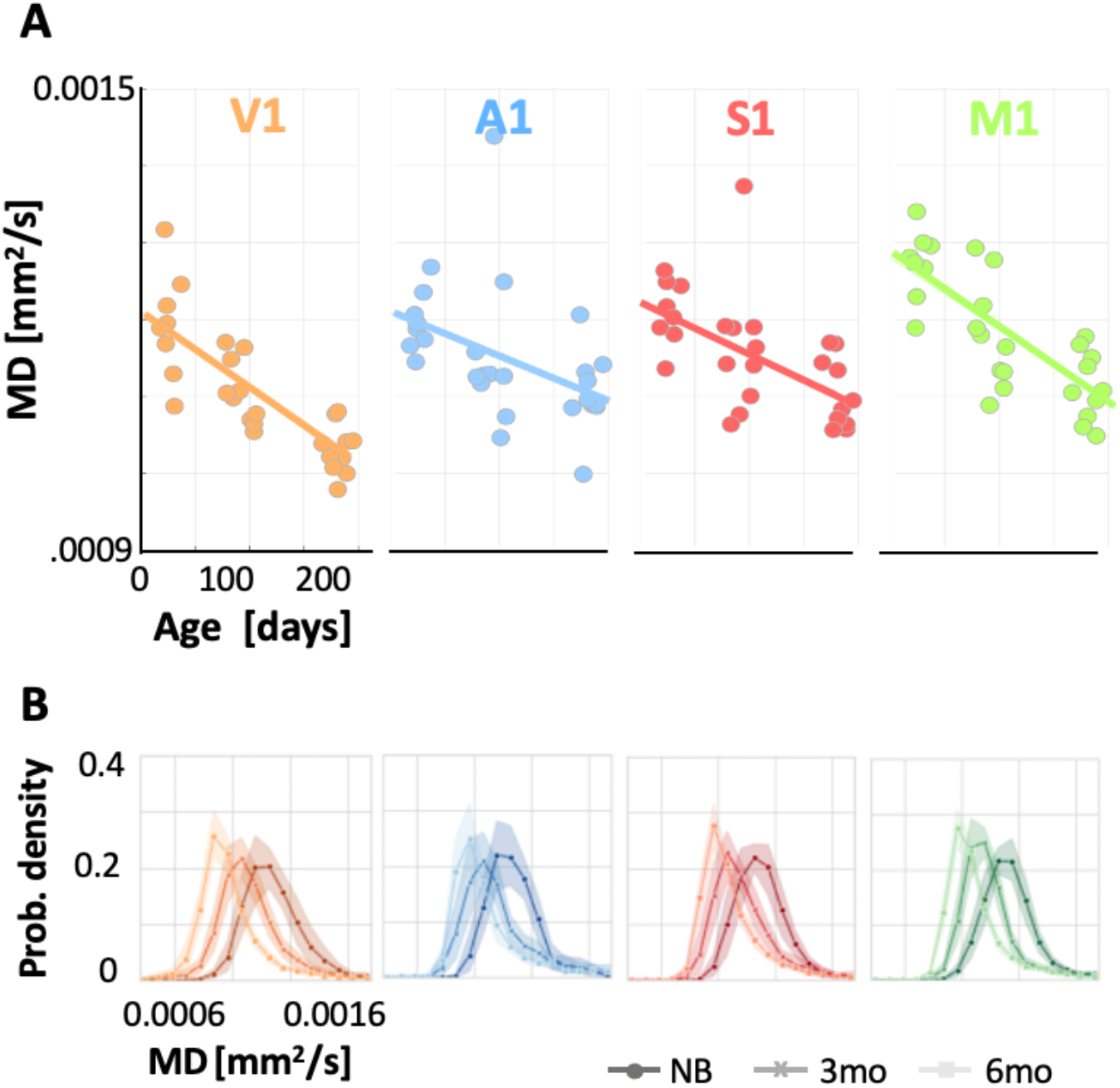
Mean MD in four early sensory regions in the left hemispheres decreases with age in the first six months of infant life. (A) Significant decrease in MD with age in the four primary sensory-motor cortices (*V1: primary visual area, A1: primary auditory area, S1: primary somatosensory area, M1: primary motor area)*. Slopes and p-values in **Supplementary Table S3**. Each dot represents the mean MD of an infant’s region of interest. (B) Distribution of MD across voxels of the four primary sensory-motor cortices shows broader distribution of MD at birth as compared to that at six months (darker colored lines: newborns). In all primary sensory-motor regions the mean MD significantly decreases from 0.0012mm^2^/s ±6.8×10-5 (M±SD) at 0 month to 0.0011 mm^2^/s ± 7.59×10-5 at 3 months to 0.0011 ± 5.52×10-5 at 6 months. Solid lines indicate mean, shaded region indicates standard error across 10 participants at each timepoint *NB:* newborns; *3 mo:* 3 month-olds; *6 mo:* 6 month-olds.

**Supplementary Figure 5.**
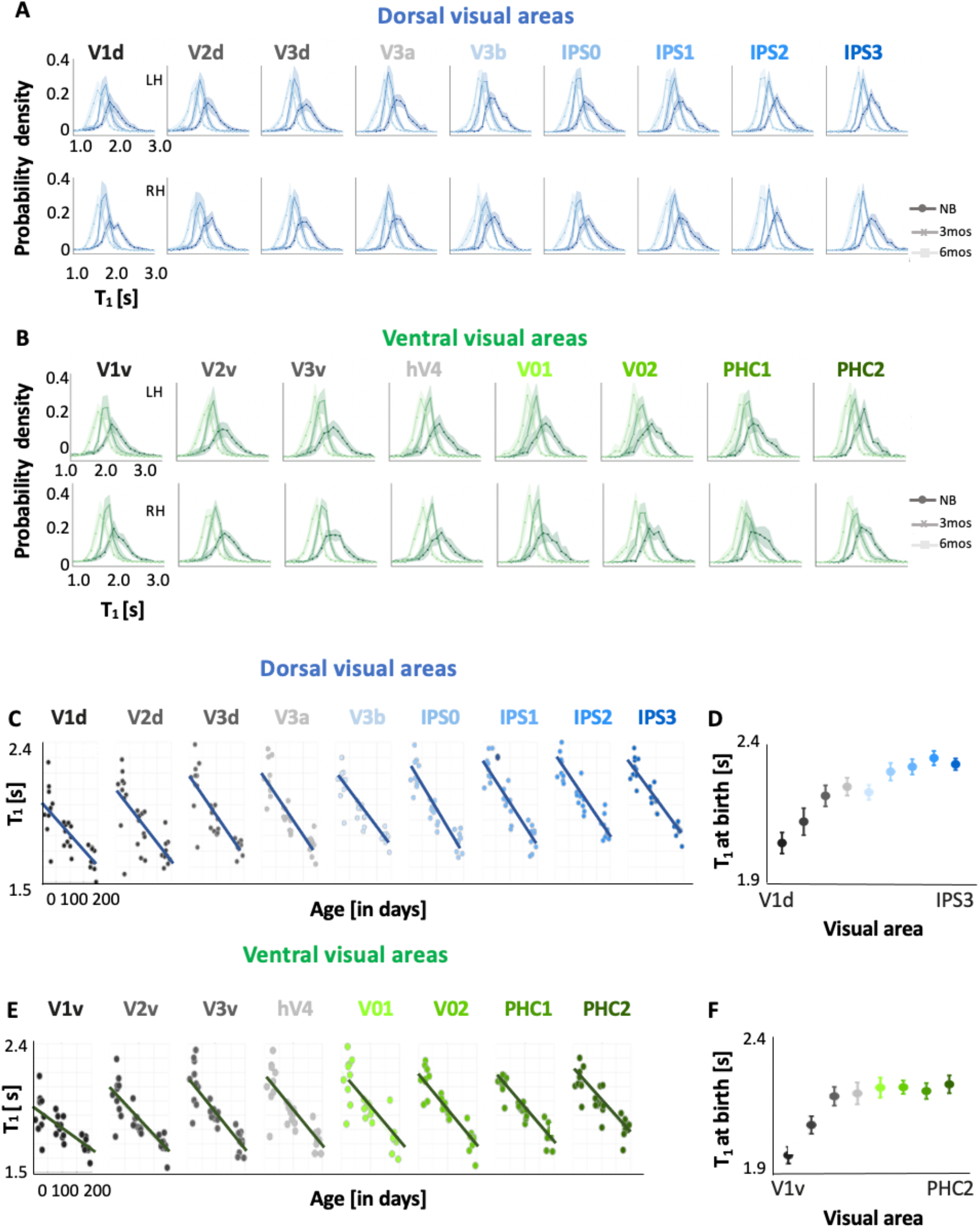
Hierarchical T_1_ development in the left dorsal and ventral visual streams during early infancy. (A, B)Distributions of T_1_ in dorsal (A) and ventral (B) visual regions in the left and right hemispheres shift leftward with age in the first six months of infant life. *LH/RH:* left/right hemisphere. (C, E) Developmental trajectory of T_1_ in the first 6 months of life (C) Left dorsal stream (V1d to IPS3). (E) left ventral stream (V1v to PHC2). T_1_ linearly decreases in all ventral and dorsal visual areas. Each dot represents mean T_1_ per ROI per infant. (D,E) Mean T_1_ at birth in the left dorsal (D) and ventral (E) streams (measured as intercepts of linear mixed model per region) shows a hierarchical development with lower T_1_ in earlier visual areas of theses stream. Comparison of LMM estimates of T_1_ at birth (intercept) across visual areas shows a gradual increase of cortical T_1_ at birth across the visual hierarchy, in both streams.

**Supplementary Figure 6.**
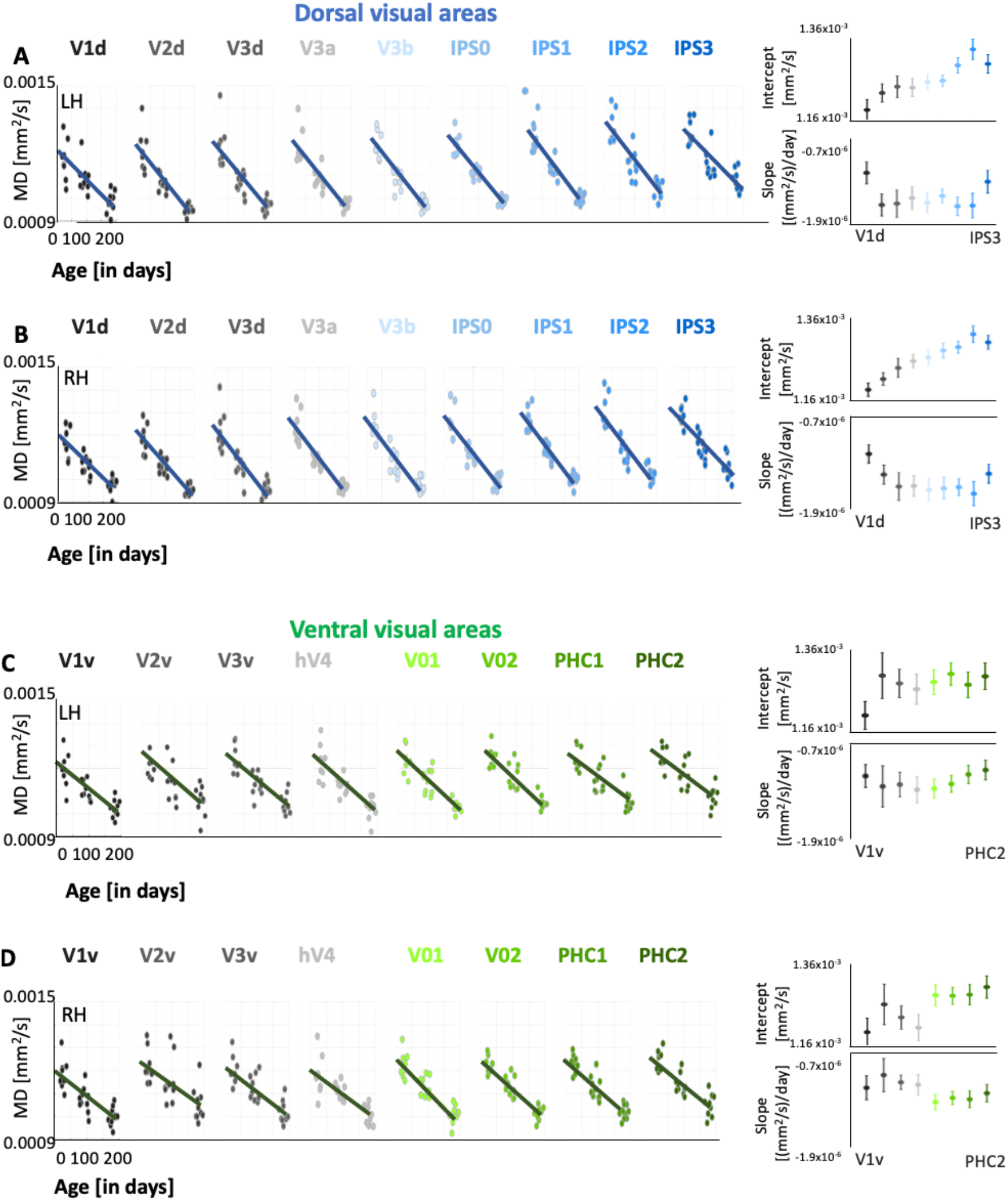
Hierarchical MD development in the dorsal and ventral visual streams in the left and right hemisphere during early infancy. (A,B). Left to right shows the developmental trajectory of MD in the first 6 months of life, in the left (A) and right (B) dorsal visual streams (V1d to IPS3) respectively. MD linearly decreases in all dorsal visual areas; slopes and p-values in **Supplementary Table S5**. Each dot represents mean MD per ROI per infant. The slopes and intercepts of each LMM are shown on the right-side of each panel. Mean MD at birth in dorsal (measured as intercepts of linear mixed model per region) shows a hierarchical development with lower MD in posterior than dorsal/anterior regions of the stream. (C,D) same as in A,B but for ventral stream visual areas (V1v to PHC2). Interestingly, comparison of LMM estimates of MD at birth (intercept) across visual areas shows a gradual increase of cortical MD at birth across the visual hierarchy, in both the dorsal visual stream, from V1d [left: 0.0012mm^2^/s±2.21×10-5; right 0.0012mm^2^/s±1.63×10-5] to IPS3 [left: 0.0013mm^2^/s±2.205×10-5; right: 0.0013mm^2^/s±1.70×10-5] and the ventral visual stream, from V1v [left: 0.0012mm^2^/s ±1.83×10-5; right:0.0012mm^2^/s±1.97×10-5] to PHC2 [left: 0.013mm^2^/s +1.74×10-5; right: 0.0013mm^2^/s ±1.58×10-5]. *LH/RH: left/right hemisphere*.

**Supplementary Figure 7.**
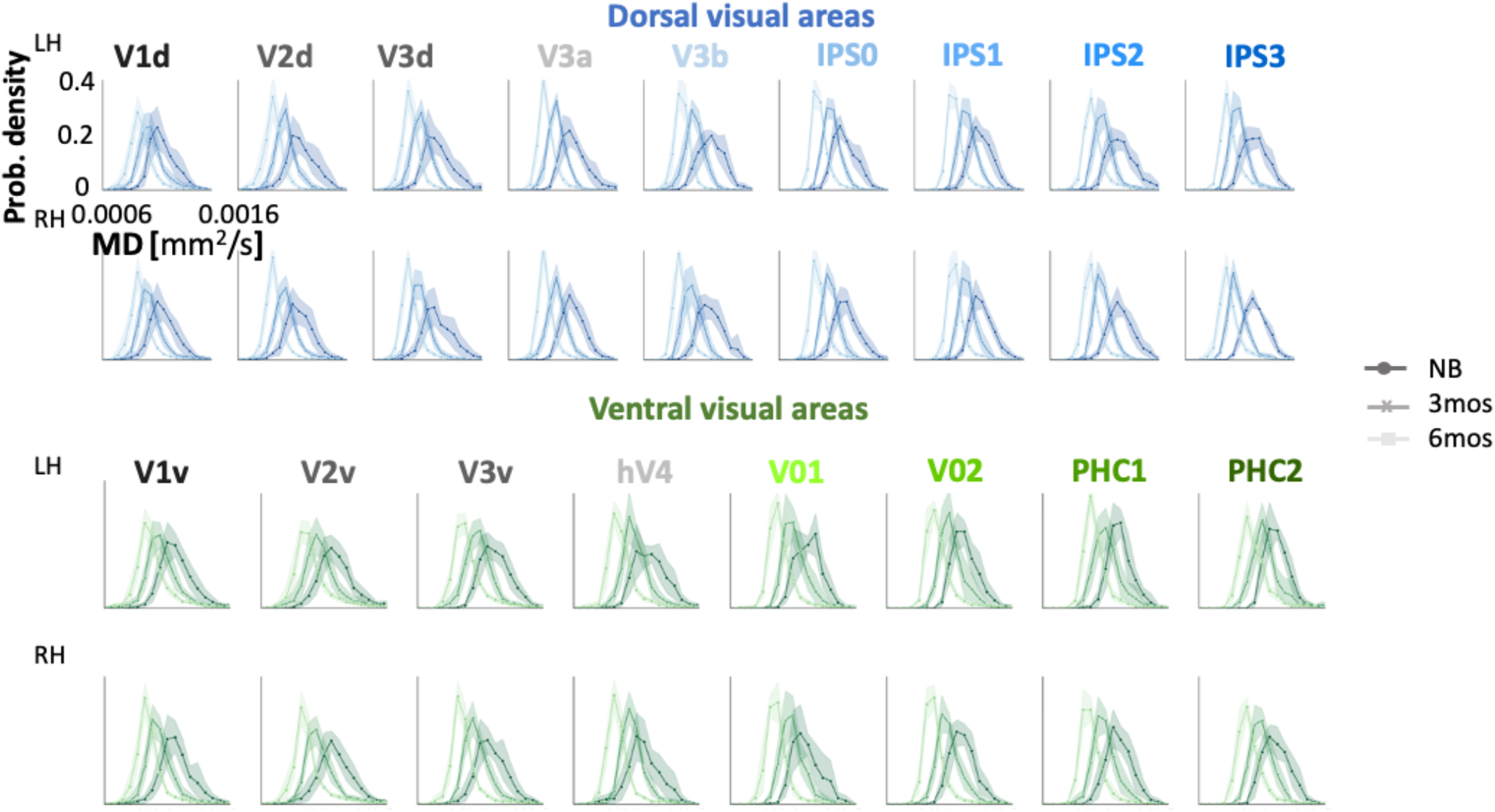
Distributions of MD in dorsal and ventral visual regions in the left and right hemisphere in newborns (NB), 3-month-olds (3mos) and 6-month-olds (6mo). Distributions shift leftward with age in the first six months of infant life indicating the MD systematically decreases. *LH/RH: left/right hemisphere. Lines: mean; Shaded areas: SE of the mean across 10 participants per time point*.

**Supplementary Figure 8.**
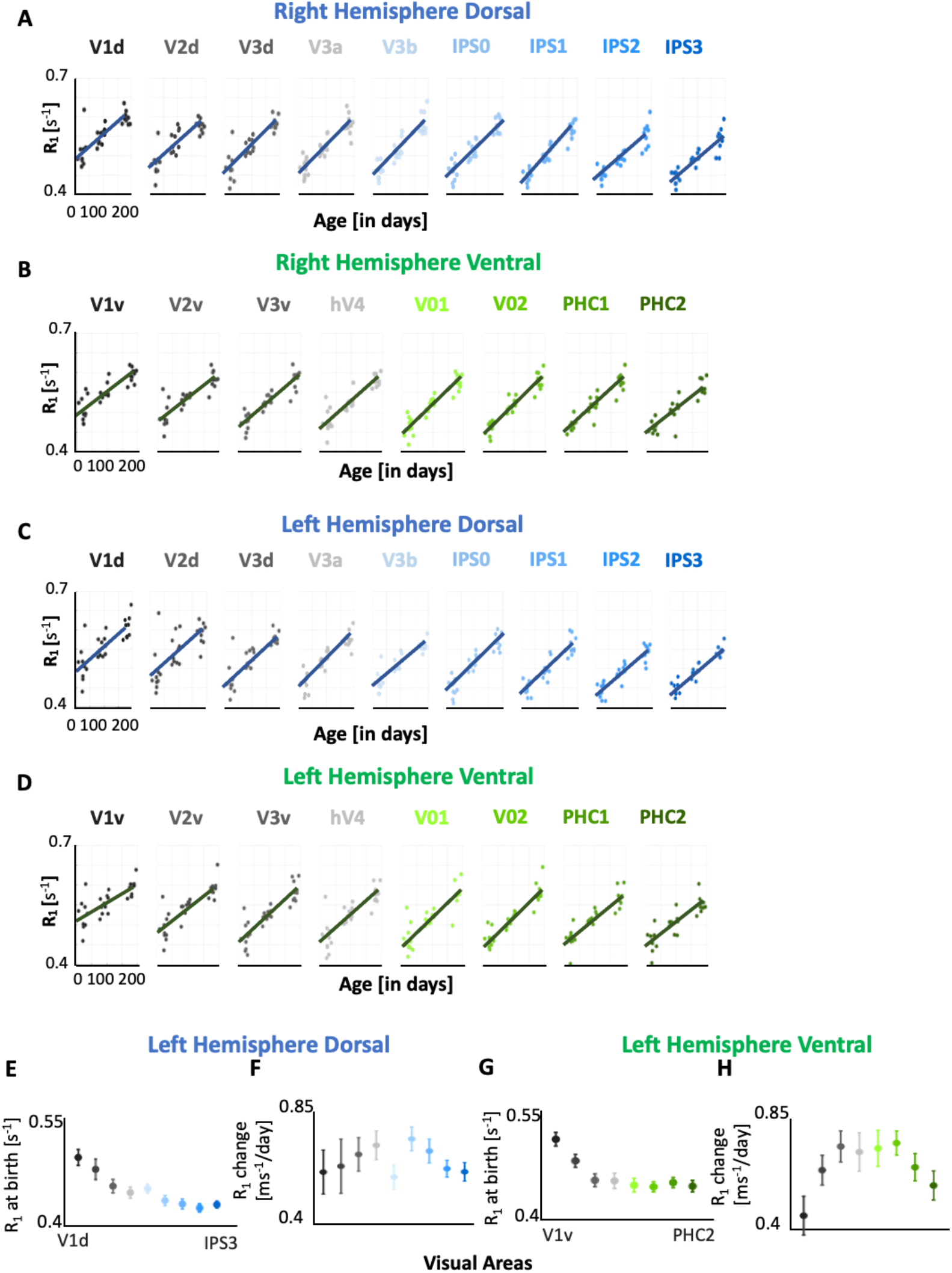
Hierarchical R_1_ development in the dorsal and ventral visual streams during early infancy. (A-D). Developmental trajectory of R_1_ in the first 6 months of life, in the right dorsal (A), right ventral (B), left dorsal (C) and left ventral (D) visual streams. R_1_ linearly increases in all visual areas. Each dot represents mean R_1_ per ROI per infant. (E-H) The intercepts and slopes of left hemisphere LMMs.

**Supplementary Table S1.**
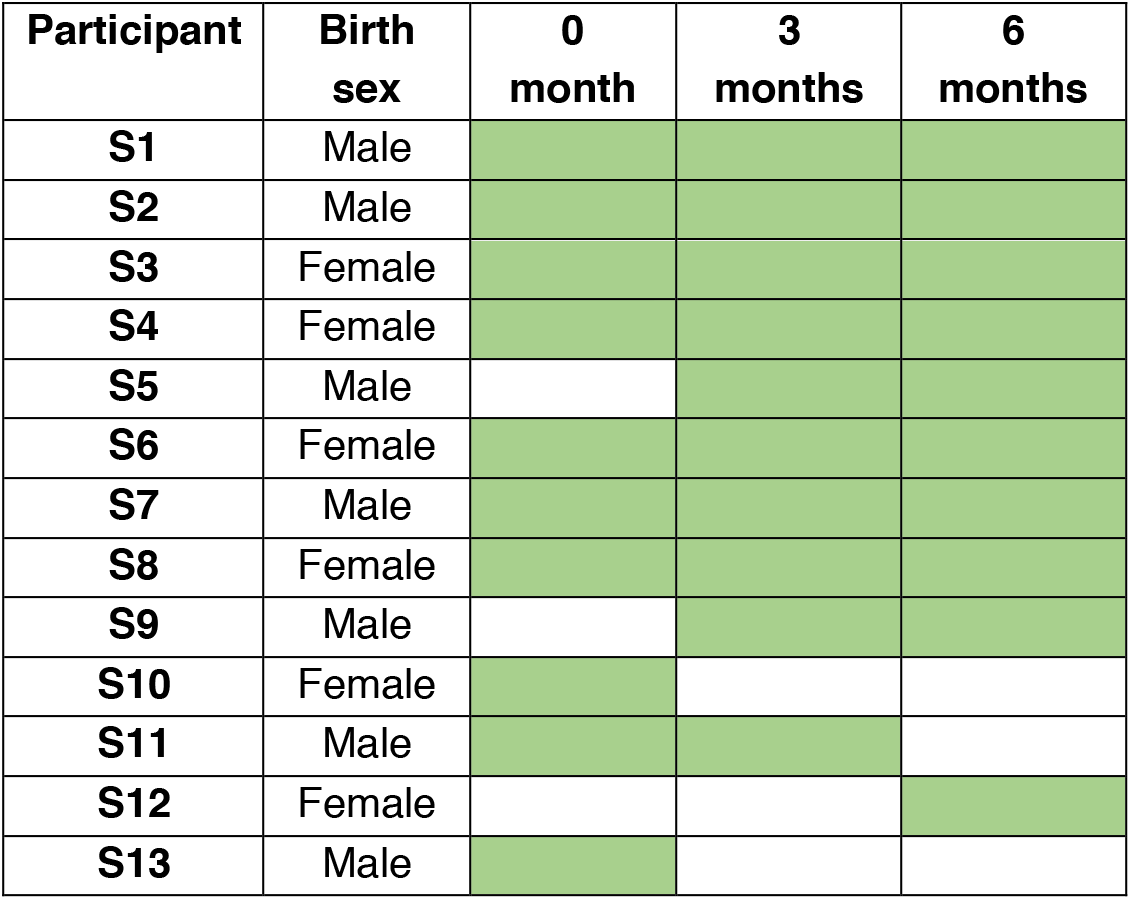
Timepoints completed by infants included in the study. *Green*: scanned, *white:* missing timepoint.

**Supplementary Table S2.** Statistical significance and parameters of linear mixed models (LMMs) quantifying the relationship between mean T_1_ and age in primary sensory cortices (related to **Fig. 1** and **Supplementary S3**). *LH/RH:* left/right hemisphere; Primary Visual (V1); Primary Auditory (A1); Primary Somatosensory (S1); Primary Motor (M1). *InC*: intercept [sec]; Slope units: [sec]/age in days. All values survive Bonferroni correction P<0.001.

| Primary<br>Sensory<br>Regions | Slope<br>(LH) | InC<br>(LH) | P-value<br>(LH) | Slope<br>(RH) | InC<br>(RH) | P-value<br>(RH) |
| --- | --- | --- | --- | --- | --- | --- |
| V1 | -0.0019 | 2.01 | $1.48 \times 10^{-9}$ | -0.0020 | 2.03 | $2.22 \times 10^{-11}$ |
| A1 | -0.0013 | 1.99 | $1.33 \times 10^{-7}$ | -0.0017 | 2.04 | $2.63 \times 10^{-9}$ |
| S1 | -0.0018 | 2.09 | $2.44 \times 10^{-12}$ | -0.0020 | 2.11 | $9.67 \times 10^{-12}$ |
| M1 | -0.0019 | 2.08 | $8.86 \times 10^{-13}$ | -0.0019 | 2.10 | $3.73 \times 10^{-10}$ |

**Supplementary Table S3.** Statistical significance and parameters of LMMs quantifying the relationship between mean MD and age in primary sensory-motor cortices (related to **Supplementary Fig. S4**). *LH/RH*: left/right hemisphere; Primary Visual (V1); Primary Auditory (A1); Primary Somatosensory (S1); Primary Motor (M1). *InC:* intercept [mm^2^/sec]; Slope units: [mm^2^/sec]/age in days. All values survive Bonferroni correction P<0.001, except those in gray.

| Primary<br>Sensory<br>Regions | Slope<br>(LH) | InC<br>(LH) | P-value<br>(LH) | Slope<br>(RH) | InC<br>(RH) | P-value<br>(RH) |
| --- | --- | --- | --- | --- | --- | --- |
| V1 | -1.01x10<sup>-6</sup> | 0.00121 | 6.13x10<sup>-8</sup> | -1.08x10<sup>-6</sup> | 0.0012123 | 9.86x10<sup>-10</sup> |
| A1 | -5.70x10<sup>-7</sup> | 0.0012089 | 0.01 | -7.09x10<sup>-7</sup> | 0.0012148 | 0.002 |
| S1 | -6.40x10<sup>-7</sup> | 0.0012216 | 0.001 | -5.48x10<sup>-7</sup> | 0.0012003 | 0.0007 |
| M1 | -9.36x10<sup>-7</sup> | 0.0012801 | 8.26x10<sup>-7</sup> | -1.06x10<sup>-6</sup> | 0.0012974 | 2.59x10<sup>-5</sup> |

**Supplementary Table S4.** Statistical significance and parameters of LMMs quantifying the relationship between mean T_1_ and age in dorsal and ventral visual areas (related to **Fig. 2** and **Supplementary Fig. S5**). *LH/RH*: left/right hemisphere; *InC:* intercept [sec]; Slope units: [sec]/age in days. All values survive Bonferroni correction P<0.001.

| **Dorsal Regions** | **Slope (LH)** | **InC (LH)** | **P-value (LH)** | **Slope (RH)** | **InC (RH)** | **P-value (RH)** |
| --- | --- | --- | --- | --- | --- | --- |
| V1d | -0.0020 | 2.01 | 5.70x10<sup>-7</sup> | -0.0020 | 2.01 | 2.80x10<sup>-8</sup> |
| V2d | -0.0023 | 2.08 | 7.45x10<sup>-6</sup> | -0.0023 | 2.12 | 2.66x10<sup>-8</sup> |
| V3d | -0.0026 | 2.17 | 1.08x10<sup>-8</sup> | -0.0027 | 2.18 | 1.94x10<sup>-9</sup> |
| V3a | -0.0027 | 2.20 | 9.26x10<sup>-11</sup> | -0.0027 | 2.18 | 1.00x10<sup>-10</sup> |
| V3b | -0.0023 | 2.18 | 9.14x10<sup>-11</sup> | -0.0029 | 2.23 | 8.68x10<sup>-13</sup> |
| IPS0 | -0.0029 | 2.26 | 1.95x10<sup>-11</sup> | -0.0029 | 2.23 | 1.31x10<sup>-11</sup> |
| IPS1 | -0.0028 | 2.27 | 7.79x10<sup>-12</sup> | -0.0030 | 2.29 | 3.58x10<sup>-15</sup> |
| IPS2 | -0.0026 | 2.30 | 1.71x10<sup>-11</sup> | -0.0025 | 2.27 | 1.90x10<sup>-12</sup> |
| IPS3 | -0.0025 | 2.28 | 1.68x10<sup>-14</sup> | -0.0025 | 2.29 | 3.34x10<sup>-11</sup> |

| **Ventral Regions** | **Slope (LH)** | **InC (LH)** | **P-value (LH)** | **Slope (RH)** | **InC (RH)** | **P-value (RH)** |
| --- | --- | --- | --- | --- | --- | --- |
| V1v | -0.0015 | 1.96 | 3.75x10<sup>-6</sup> | -0.0019 | 2.01 | 8.11x10<sup>-9</sup> |
| V2v | -0.0022 | 2.07 | 1.25x10<sup>-8</sup> | -0.0021 | 2.07 | 8.06x10<sup>-9</sup> |
| V3v | -0.0026 | 2.17 | 1.53x10<sup>-9</sup> | -0.0024 | 2.14 | 2.02x10<sup>-11</sup> |
| hV4 | -0.0026 | 2.18 | 5.02x10<sup>-8</sup> | -0.0024 | 2.15 | 2.86x10<sup>-10</sup> |
| V01 | -0.0027 | 2.20 | 4.81x10<sup>-9</sup> | -0.0027 | 2.21 | 2.28x10<sup>-12</sup> |
| V02 | -0.0027 | 2.21 | 4.09x10<sup>-13</sup> | -0.0028 | 2.22 | 8.50x10<sup>-15</sup> |
| PHC1 | -0.0025 | 2.19 | 1.59x10<sup>-10</sup> | -0.0026 | 2.19 | 2.28x10<sup>-13</sup> |
| PHC2 | -0.0023 | 2.22 | 1.49x10<sup>-8</sup> | -0.0025 | 2.21 | 3.62x10<sup>-10</sup> |

**Supplementary Table S5.**
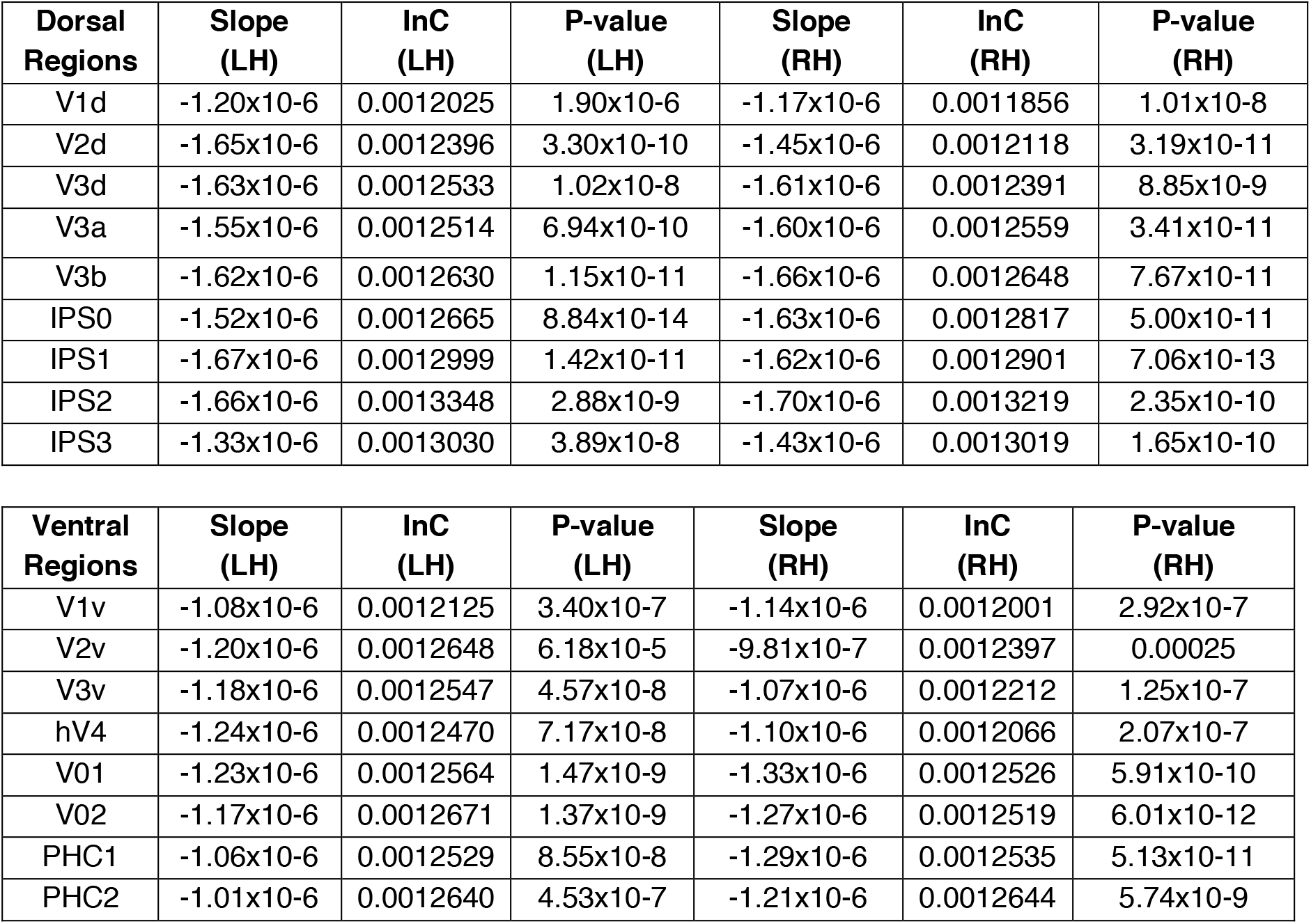
Statistical significance and parameters of LMMs quantifying the relationship between mean MD and age in dorsal and ventral visual areas (related to **Fig. 2** and **Supplementary Fig. S6**). *LH/RH*: left/right hemisphere; *InC*: Intercept [mm^2^/sec]; Slope units: [mm^2^/sec]/age in days. All values survive Bonferroni correction P<0.001.

| **Dorsal Regions** | **Slope (LH)** | **InC (LH)** | **P-value (LH)** | **Slope (RH)** | **InC (RH)** | **P-value (RH)** |
| --- | --- | --- | --- | --- | --- | --- |
| V1d | -1.20x10<sup>-6</sup> | 0.0012025 | 1.90x10<sup>-6</sup> | -1.17x10<sup>-6</sup> | 0.0011856 | 1.01x10<sup>-8</sup> |
| V2d | -1.65x10<sup>-6</sup> | 0.0012396 | 3.30x10<sup>-10</sup> | -1.45x10<sup>-6</sup> | 0.0012118 | 3.19x10<sup>-11</sup> |
| V3d | -1.63x10<sup>-6</sup> | 0.0012533 | 1.02x10<sup>-8</sup> | -1.61x10<sup>-6</sup> | 0.0012391 | 8.85x10<sup>-9</sup> |
| V3a | -1.55x10<sup>-6</sup> | 0.0012514 | 6.94x10<sup>-10</sup> | -1.60x10<sup>-6</sup> | 0.0012559 | 3.41x10<sup>-11</sup> |
| V3b | -1.62x10<sup>-6</sup> | 0.0012630 | 1.15x10<sup>-11</sup> | -1.66x10<sup>-6</sup> | 0.0012648 | 7.67x10<sup>-11</sup> |
| IPS0 | -1.52x10<sup>-6</sup> | 0.0012665 | 8.84x10<sup>-14</sup> | -1.63x10<sup>-6</sup> | 0.0012817 | 5.00x10<sup>-11</sup> |
| IPS1 | -1.67x10<sup>-6</sup> | 0.0012999 | 1.42x10<sup>-11</sup> | -1.62x10<sup>-6</sup> | 0.0012901 | 7.06x10<sup>-13</sup> |
| IPS2 | -1.66x10<sup>-6</sup> | 0.0013348 | 2.88x10<sup>-9</sup> | -1.70x10<sup>-6</sup> | 0.0013219 | 2.35x10<sup>-10</sup> |
| IPS3 | -1.33x10<sup>-6</sup> | 0.0013030 | 3.89x10<sup>-8</sup> | -1.43x10<sup>-6</sup> | 0.0013019 | 1.65x10<sup>-10</sup> |

| **Ventral Regions** | **Slope (LH)** | **InC (LH)** | **P-value (LH)** | **Slope (RH)** | **InC (RH)** | **P-value (RH)** |
| --- | --- | --- | --- | --- | --- | --- |
| V1v | -1.08x10<sup>-6</sup> | 0.0012125 | 3.40x10<sup>-7</sup> | -1.14x10<sup>-6</sup> | 0.0012001 | 2.92x10<sup>-7</sup> |
| V2v | -1.20x10<sup>-6</sup> | 0.0012648 | 6.18x10<sup>-5</sup> | -9.81x10<sup>-7</sup> | 0.0012397 | 0.00025 |
| V3v | -1.18x10<sup>-6</sup> | 0.0012547 | 4.57x10<sup>-8</sup> | -1.07x10<sup>-6</sup> | 0.0012212 | 1.25x10<sup>-7</sup> |
| hV4 | -1.24x10<sup>-6</sup> | 0.0012470 | 7.17x10<sup>-8</sup> | -1.10x10<sup>-6</sup> | 0.0012066 | 2.07x10<sup>-7</sup> |
| V01 | -1.23x10<sup>-6</sup> | 0.0012564 | 1.47x10<sup>-9</sup> | -1.33x10<sup>-6</sup> | 0.0012526 | 5.91x10<sup>-10</sup> |
| V02 | -1.17x10<sup>-6</sup> | 0.0012671 | 1.37x10<sup>-9</sup> | -1.27x10<sup>-6</sup> | 0.0012519 | 6.01x10<sup>-12</sup> |
| PHC1 | -1.06x10<sup>-6</sup> | 0.0012529 | 8.55x10<sup>-8</sup> | -1.29x10<sup>-6</sup> | 0.0012535 | 5.13x10<sup>-11</sup> |
| PHC2 | -1.01x10<sup>-6</sup> | 0.0012640 | 4.53x10<sup>-7</sup> | -1.21x10<sup>-6</sup> | 0.0012644 | 5.74x10<sup>-9</sup> |

**Supplementary Table S6.** **Demographic information of postmortem human brain tissue samples used for the transcriptomic gene analysis**. The data is from the Brain Span Atlas portal at www.brainspan.org. Developmental stages from mid to late prenatal to early infancy were used for the differential gene analysis. *PCW:* post conceptual weeks, *M:* months; *F/M*: female/male; *E* = European, *A* = African American, *H* = Hispanic.

| Subject ID | Age at Death | Gender | Ethnicity |
| --- | --- | --- | --- |
|  | Prenatal samples (in PCW) |  |  |
| H376.IV.53 | 19 | F | H |
| H376.IV.51 | 21 | F | E |
| H376.IV.54 | 21 | M | A |
| H376.IV.50 | 22 | M | E |
| H376.V.51 | 25 | F | H |
| H376.V.52 | 26 | F | A |
| H376.V.53 | 37 | M | E |
|  | Postnatal samples (in M) |  |  |
| H376.VI.50 | 4 | M | E |
| H376.VI.51 | 4 | M | E |
| H376.VI.52 | 4 | M | A |

**Supplementary Table S7.** Brain regions used for the transcriptomic gene analysis. Tissue samples from prenatal and postnatal human primary sensory-motor, temporal, and parietal cortices profiled by RNA sequencing. Primary motor cortex (M1, BA4), Primary visual cortex (V1, BA17), Primary somatosensory cortex (S1, BA1-3), Primary auditory temporal cortex (A1, BA41), Posterior inferior parietal cortex (IPC, BA40), Posterior superior temporal cortex (STC, BA22), Inferior temporal cortex (ITC, BA20); *PCW*: post conceptual weeks, *M:* months.

| Subject ID | Age | Regions |
| --- | --- | --- |
| H376.IV.53 | 19 pcw | M1C-S1C |
| H376.IV.53 | 19 pcw | IPC |
| H376.IV.53 | 19 pcw | A1C |
| H376.IV.53 | 19 pcw | STC |
| H376.IV.53 | 19 pcw | V1C |
| H376.IV.51 | 21 pcw | ITC |
| H376.IV.54 | 21 pcw | M1C |
| H376.IV.54 | 21 pcw | S1C |
| H376.IV.54 | 21 pcw | IPC |
| H376.IV.54 | 21 pcw | STC |
| H376.IV.54 | 21 pcw | ITC |
| H376.IV.54 | 21 pcw | V1C |
| H376.IV.50 | 22 pcw | M1C |
| H376.IV.50 | 22 pcw | S1C |
| H376.IV.50 | 22 pcw | IPC |
| H376.IV.50 | 22 pcw | A1C |
| H376.IV.50 | 22 pcw | STC |
| H376.IV.50 | 22 pcw | ITC |
| H376.IV.50 | 22 pcw | V1C |
| H376.V.51 | 25 pcw | A1C |
| H376.V.52 | 26 pcw | V1C |
| H376.V.52 | 26 pcw | STC |
| H376.V.53 | 37 pcw | M1C |
| H376.V.53 | 37 pcw | S1C |
| H376.V.53 | 37 pcw | IPC |
| H376.V.53 | 37 pcw | A1C |
| H376.V.53 | 37 pcw | STC |
| H376.V.53 | 37 pcw | ITC |
| H376.V.53 | 37 pcw | V1C |
| H376.VI.50 | 4 M | ITC |
| H376.VI.50 | 4 M | STC |
| H376.VI.50 | 4 M | V1C |
| H376.VI.51 | 4 M | M1C |
| H376.VI.51 | 4 M | A1C |
| H376.VI.51 | 4 M | STC |
| H376.VI.51 | 4 M | ITC |
| H376.VI.52 | 4 M | M1C |
| H376.VI.52 | 4 M | S1C |
| H376.VI.52 | 4 M | IPC |
| H376.VI.52 | 4 M | A1C |
| H376.VI.52 | 4 M | STC |
| H376.VI.52 | 4 M | V1C |
| H376.VI.52 | 4 M | ITC |

**Supplementary Table S8.** The gene ontology (GO) list related to Figure 3C. This GO list includes the information on the biological processes related to the 95 most differentially expressed genes, listed by statistical significance of functional enrichment analysis. Complete gene ontology lists without and with background gene sets can be found on GitHub: https://github.com/VPNL/babies_graymatter/genes/Dataset1 and https://github.com/VPNL/babies_graymatter/genes/Dataset2

| Category | ID | Name | p-value<br>FDR |
| --- | --- | --- | --- |
| GO: Molecular Function | GO:0019911 | structural constituent of myelin sheath | 2.05E-05 |
| GO: Molecular Function | GO:0042165 | neurotransmitter binding | 5.57E-04 |
| GO: Biological Process | GO:0098916 | anterograde trans-synaptic signaling | 1.29E-12 |
| GO: Biological Process | GO:0007268 | chemical synaptic transmission | 1.29E-12 |
| GO: Biological Process | GO:0099537 | trans-synaptic signaling | 1.29E-12 |
| GO: Biological Process | GO:0099536 | synaptic signaling | 1.34E-12 |
| GO: Biological Process | GO:0007267 | cell-cell signaling | 5.19E-12 |
| GO: Biological Process | GO:0050804 | modulation of chemical synaptic transmission | 5.82E-08 |
| GO: Biological Process | GO:0099177 | regulation of trans-synaptic signaling | 5.82E-08 |
| GO: Biological Process | GO:0048167 | regulation of synaptic plasticity | 2.89E-05 |
| GO: Biological Process | GO:0048666 | neuron development | 3.56E-05 |
| GO: Biological Process | GO:1900449 | regulation of glutamate receptor signaling pathway | 3.56E-05 |
| GO: Biological Process | GO:0030030 | cell projection organization | 6.34E-05 |
| GO: Biological Process | GO:0030182 | neuron differentiation | 1.02E-04 |
| GO: Biological Process | GO:0042552 | myelination | 1.38E-04 |
| GO: Biological Process | GO:0032291 | axon ensheathment in central nervous system | 1.38E-04 |
| GO: Biological Process | GO:0022010 | central nervous system myelination | 1.38E-04 |
| GO: Biological Process | GO:0008366 | axon ensheathment | 1.38E-04 |
| GO: Biological Process | GO:0007272 | ensheathment of neurons | 1.38E-04 |
| GO: Cellular Component | GO:0043005 | neuron projection | 2.98E-15 |
| GO: Cellular Component | GO:0030424 | axon | 2.10E-14 |
| GO: Cellular Component | GO:0036477 | somatodendritic compartment | 1.16E-11 |
| GO: Cellular Component | GO:0045202 | synapse | 9.90E-10 |
| GO: Cellular Component | GO:0043025 | neuronal cell body | 1.23E-09 |
| GO: Cellular Component | GO:0044297 | cell body | 7.07E-09 |
| GO: Cellular Component | GO:0098794 | postsynapse | 3.81E-07 |
| GO: Cellular Component | GO:0044304 | main axon | 1.37E-06 |
| GO: Cellular Component | GO:0098793 | presynapse | 1.02E-05 |
| GO: Cellular Component | GO:0097060 | synaptic membrane | 2.76E-05 |
| GO: Cellular Component | GO:0030425 | dendrite | 2.78E-05 |
| GO: Cellular Component | GO:0097447 | dendritic tree | 2.78E-05 |

**Supplementary Table S9.** Statistical significance and parameters of LMMs quantifying the relationship between mean R_1_ and age in dorsal and ventral visual areas (related to **Fig. 4** and **Supplementary Fig. S8**). *LH/RH:* left/right hemisphere; *Inc:* Intercept [s^−1^]. Slope units: [msec^−1^]/age in days. All values survive Bonferroni correction P<0.001.

| Dorsal Regions | Slope (LH) | Inc (LH) | P-value (LH) | Slope (RH) | Inc (RH) | P-value (RH) |
| --- | --- | --- | --- | --- | --- | --- |
| V1d | 0.6166 | 0.49 | $3.02 \times 10^{-07}$ | 0.5903 | 0.49 | $3.78 \times 10^{-08}$ |
| V2d | 0.6396 | 0.48 | $1.59 \times 10^{-05}$ | 0.6254 | 0.46 | $9.52 \times 10^{-09}$ |
| V3d | 0.6858 | 0.45 | $3.34 \times 10^{-09}$ | 0.7259 | 0.45 | $2.06 \times 10^{-10}$ |
| V3a | 0.7216 | 0.44 | $8.07 \times 10^{-12}$ | 0.7283 | 0.45 | $5.04 \times 10^{-11}$ |
| V3b | 0.5970 | 0.45 | $1.43 \times 10^{-11}$ | 0.7843 | 0.44 | $1.26 \times 10^{-12}$ |
| IPS0 | 0.7466 | 0.43 | $1.21 \times 10^{-12}$ | 0.7636 | 0.44 | $1.14 \times 10^{-12}$ |
| IPS1 | 0.6990 | 0.43 | $5.69 \times 10^{-12}$ | 0.7707 | 0.42 | $8.66 \times 10^{-16}$ |
| IPS2 | 0.6298 | 0.42 | $3.26 \times 10^{-12}$ | 0.6411 | 0.43 | $5.35 \times 10^{-12}$ |
| IPS3 | 0.6176 | 0.43 | $2.56 \times 10^{-15}$ | 0.6196 | 0.43 | $2.29 \times 10^{-11}$ |

| Ventral Regions | Slope (LH) | Inc (LH) | P-value (LH) | Slope (RH) | Inc (RH) | P-value (RH) |
| --- | --- | --- | --- | --- | --- | --- |
| V1v | 0.4576 | 0.50 | $2.24 \times 10^{-06}$ | 0.5557 | 0.49 | $2.45 \times 10^{-09}$ |
| V2v | 0.6304 | 0.48 | $2.72 \times 10^{-09}$ | 0.5889 | 0.47 | $2.29 \times 10^{-09}$ |
| V3v | 0.7204 | 0.45 | $2.52 \times 10^{-10}$ | 0.6689 | 0.46 | $4.43 \times 10^{-12}$ |
| hV4 | 0.6994 | 0.45 | $1.46 \times 10^{-08}$ | 0.6573 | 0.46 | $3.24 \times 10^{-11}$ |
| V01 | 0.7131 | 0.44 | $1.46 \times 10^{-09}$ | 0.7281 | 0.44 | $1.92 \times 10^{-13}$ |
| V02 | 0.7334 | 0.44 | $4.09 \times 10^{-13}$ | 0.7644 | 0.44 | $1.96 \times 10^{-15}$ |
| PHC1 | 0.6415 | 0.45 | $9.08 \times 10^{-12}$ | 0.6999 | 0.45 | $1.54 \times 10^{-13}$ |
| PHC2 | 0.5722 | 0.44 | $4.25 \times 10^{-09}$ | 0.6540 | 0.44 | $1.17 \times 10^{-11}$ |

